# Chipper: Open-source software for semi-automated segmentation and analysis of birdsong and other natural sounds

**DOI:** 10.1101/807974

**Authors:** Abigail M. Searfoss, James C. Pino, Nicole Creanza

## Abstract

1. Audio recording devices have changed significantly over the last 50 years, making large datasets of recordings of natural sounds, such as birdsong, easier to obtain. This increase in digital recordings necessitates an increase in high-throughput methods of analysis for researchers. Specifically, there is a need in the community for open-source methods that are tailored to recordings of varying qualities and from multiple species collected in nature.
2. We developed Chipper, a Python-based software to semi-automate both the segmentation of acoustic signals and the subsequent analysis of their frequencies and durations. For avian recordings, we provide widgets to best determine appropriate thresholds for noise and syllable similarity, which aid in calculating note measurements and determining syntax. In addition, we generated a set of synthetic songs with various levels of background noise to test Chipper’s accuracy, repeatability, and reproducibility.
3. Chipper provides an effective way to quickly generate reproducible estimates of birdsong features. The cross-platform graphical user interface allows the user to adjust parameters and visualize the resulting spectrogram and signal segmentation, providing a simplified method for analyzing field recordings.
4. Chipper streamlines the processing of audio recordings with multiple user-friendly tools and is optimized for multiple species and varying recording qualities. Ultimately, Chipper supports the use of citizen-science data and increases the feasibility of large-scale multi-species birdsong studies.

## Introduction

Acoustic communication is one of the few natural behaviors that can be easily recorded, digitized, and studied (Cocroft & Rodríguez 2005; Catchpole & Slater 2008; Ryan & Guerra 2014; Garland *et al*. 2017). Often, behavioral studies involve observing animals in the laboratory, which can lead to fundamental insights but can potentially alter their natural behavior (Marler & Peters 1977; Searcy 1984; Fehér *et al*. 2009). In addition, scientists can collect acoustic sounds in the wild without disturbing animals, eliminating the potential influence of the laboratory environment on the behavior of interest but limiting the types of experiments that are possible (Grant & Grant 1996; Williams *et al*. 2013; Shizuka, Ross Lein & Chilton 2016; Lachlan, Ratmann & Nowicki 2018). Moreover, recordings can be pooled across sources—professionals and hobbyists, analog and digital, old and new—providing a vast dataset that spans many years and large geographic scales (Bolus 2014; Roach & Phillmore 2017). Thus, audio recordings are an advantageous resource for broad-scale animal behavior research.

Birds, and particularly their songs, have been study systems in ecology and evolution for decades (Thorpe 1958; Marler & Tamura 1964). Historically, field studies of birdsong have provided insights into mating and territory-defense behaviors, evolutionary events such as speciation and hybridization, and environmental adaptation including anthropogenic impacts (Grant & Grant 1997; Slabbekoorn & Peet 2003; Nowicki & Searcy 2004; Mason *et al*. 2017; Snyder & Creanza 2019; Robinson, Snyder & Creanza 2019). These studies are often conducted with specific banded birds and direct recordings taken using parabolic microphones. Some song analysis software is well-suited to these studies, allowing users to visualize and manually select songs of interest from their field recordings for analysis (Cornell Laboratory of Ornithology; Burt 2001; Lachlan 2007; Boersma & Weenink 2019). On the other hand, laboratory experiments often use individual sound-attenuating recording chambers. Such experiments have greatly extended our understanding of the neurobiology of learning, stress, and development (Tchernichovski *et al*. 2001; Spencer *et al*. 2003). Alongside laboratory work, song analysis software has been developed to provide quantitative comparisons between individuals from a specific species, such as pupils and tutors in song-learning experiments (Tchernichovski *et al*. 2000; Lachlan 2007). In sum, fieldwork and laboratory experiments, particularly when paired with software, have made large contributions in understanding acoustic communication.

Concurrently, portable audio recording devices have changed significantly over the last 50 years, moving from large reel-to-reel devices to handheld digital recorders and smartphones, which has made collecting natural recordings much easier (Sullivan *et al*. 2009; Vellinga & Planqué 2015). This new technology has improved collection of both wild and laboratory recordings and led to an active worldwide community of citizen scientists who record and archive birdsong (Bonney *et al*. 2009; Sullivan *et al*. 2009; Silvertown 2009; Wood *et al*. 2011). Although there are many scientific questions that can be answered using these expanding citizen-science datasets of birdsong or other natural sounds (e.g. Xeno-Canto, eBird, Macaulay Library at the Cornell Lab of Ornithology), there is still a need for high-throughput and more automated methods of song analysis that address the varying quality and multi-species nature of citizen-science recordings. One R package, WarbleR, has made strides in this direction by facilitating the retrieval and analysis of songs from the Xeno-Canto repository (Araya-Salas & Smith-Vidaurre 2017). Existing signal processing toolboxes in Python are neither optimized for natural recordings nor user-friendly for researchers unfamiliar with computer programming. To reduce and streamline the manual work involved in processing databases of natural recordings, we developed Chipper, an open-source Python-based (v3.6.2) software with a Kivy-based (v1.10.0) graphical user interface, to semi-automate the segmentation and analysis of acoustic signals.

Chipper facilitates syllable segmentation and subsequent analysis of frequency, duration, and syntax, improving efficiency in using citizen-science recordings and increasing the feasibility of multi-species birdsong studies. Our software is open-source and easy to use, allowing seamless integration into other laboratories and STEM education programs; we successfully integrated Chipper into undergraduate courses and high-school projects. In particular, Chipper streamlines the song analysis process, eliminating the need to manually handle each song multiple times (**Figure 1**). In addition, we created a synthetic dataset of birdsong for testing acoustic software and conducted a thorough test of Chipper’s accuracy, repeatability, and reproducibility (**Supporting Information**).

**Figure 1.**
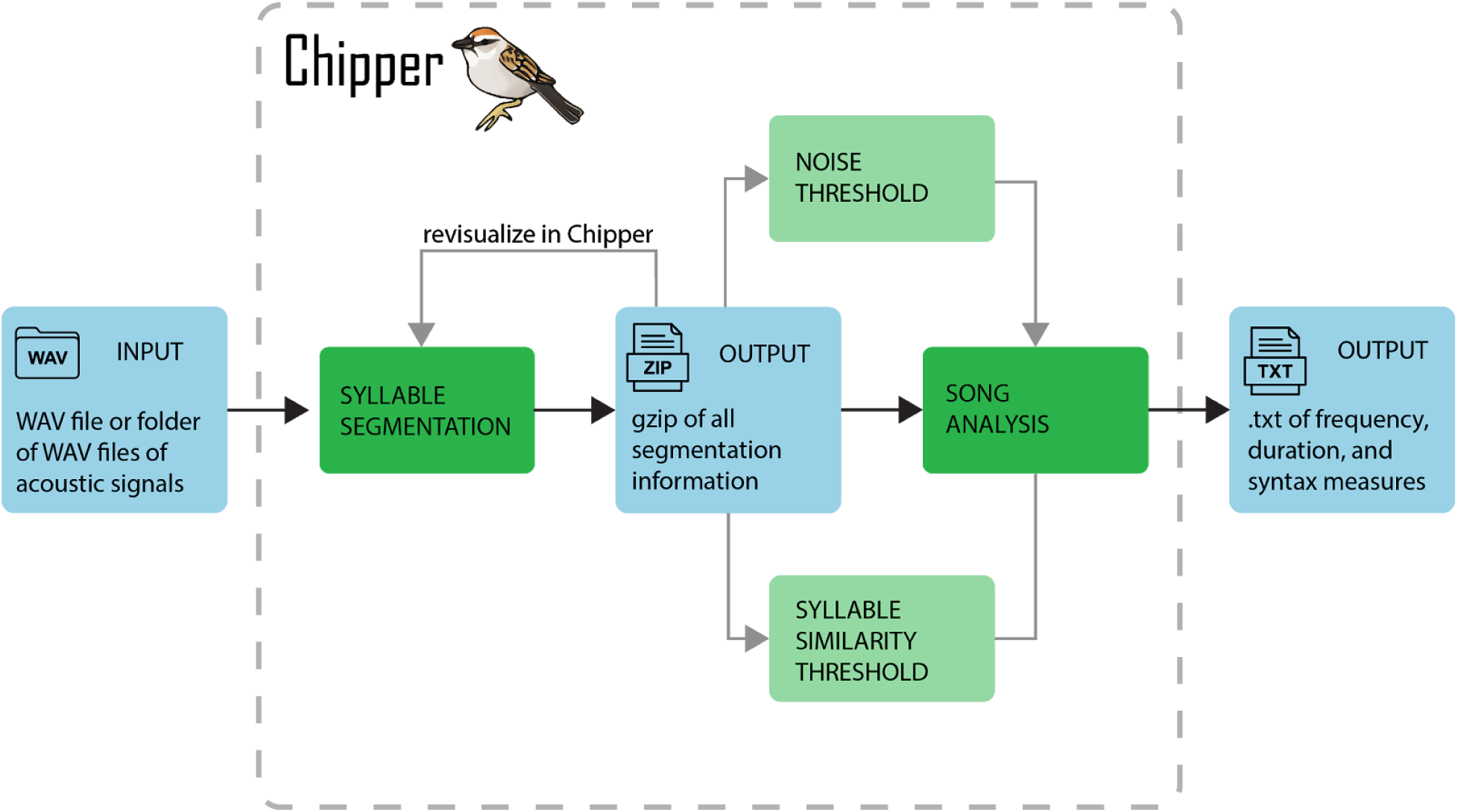
Chipper’s streamlined process of segmenting and analyzing recordings. Blue steps indicate inputs and outputs; green steps indicate Chipper widgets. Navigate through Chipper as follows: 1) Gather recordings of acoustic signals (.WAV) to input into Chipper. 2) Load files and begin with the default syllable segmentation. The user can alter segmentation parameters, viewing how this changes the quality of the signal and the segmentation. 3) Segmentation results in zipped files with all necessary information. 4) Use widgets to determine the best thresholds for noise and syllable similarity. 5) Run song analysis using these thresholds. Measurements characterizing frequencies and durations for the song, syllables, and notes are calculated. 6) Outputs all measurements into two text files.

### Chipper’s interface and capabilities

Chipper is designed to parse syllables from a bout of birdsong. We suggest recordings <3MB or ∼0.5–10 seconds, but the optimal value will differ between projects (based on sampling rate, syllable length, computing resources, etc.). For very long songs, it is advised to split the recording into multiple files before processing in Chipper. Therefore, initial selection of bouts from in-house recordings or citizen-science data (e.g. xeno-canto.org, Macaulay Library, eBird) should be completed (we suggest using Audacity), saving as WAV files. Chipper guides the user through two main steps to extract information from such WAV files: syllable segmentation and song analysis (**Figure 1**).

#### Syllable segmentation

On the Chipper landing page (**Figure 2A**), the defaults for the automated segmentation can be adjusted by the user. Next, a single WAV file or an entire folder of WAV files can be selected to begin segmentation. Chipper will then semi-automate the process of noise reduction and syllable parsing of each recorded bout of song. The syllable segmentation window (**Figure 2B**) shows two images: the top image is the spectrogram of the file and the bottom shows a binary image calculated based on user-informed parameters, with onsets (short green lines) and offsets (tall green lines) depicting the automated syllable segmentation.

**Figure 2.**
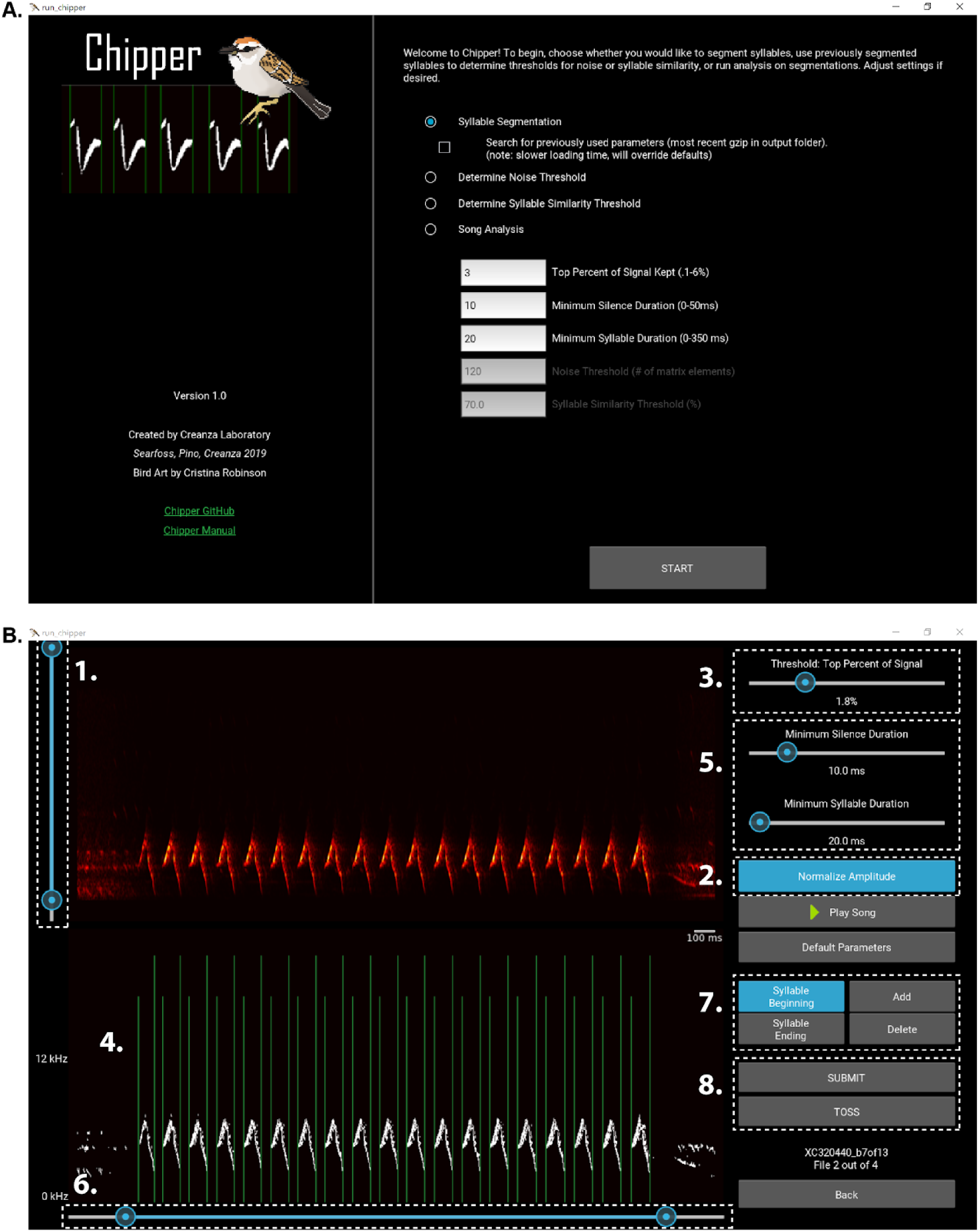
Chipper Interface. (A) Landing page of Chipper. Here the user can choose to segment songs, visualize the already segmented songs to choose thresholds for noise and syllable similarity, or run song analysis with default or user-defined thresholds. (B) Segmentation window with parameters labeled in the order that they are applied to the spectrogram and segmentation calculations (see *Syllable Segmentation* section).

The user can adjust the segmentation parameters using the sliders. With each parameter adjustment, a new binary image and corresponding onsets and offsets are calculated in the following order (numbered as in **Figure 2B**):

1. The spectrogram of the recording is created from the WAV file (method adapted from (Gardner & Magnasco 2006)), and signal outside of the two bounds of the low-pass and high-pass filter is set to zero. Colors in the resulting spectrogram are rescaled based on the remaining signal. This proves useful, as many recordings in nature have low-frequency background noise or high-frequency noise from other birds.
2. Selecting “Normalize Amplitude” rescales the amplitude across the spectrogram.
3. The “Threshold: Top Percent of Signal”, *q*, is used to find the (100-*q*)^th^ percentile of signal. Only signal above this percentile is retained and plotted in the binary image; all other signal is set to zero.
4. Syllable onsets (beginnings) and offsets (endings) are calculated by summing the columns of the spectrogram, creating a vector of total signal intensity over time. Then, the onsets are defined as the position of the first element in the matrix where signal is present after silence and the offsets as the position of the first element of the matrix with no signal after prolonged signal.
5. Two parameters act as constraints on the list of onsets and offsets—”Minimum Silence Duration” and “Minimum Syllable Duration”. If the time between the offset of one syllable and the onset of the next syllable is greater than the minimum silence duration, these boundaries are removed, combining the two syllables. Similarly, if the duration between an onset and offset of one syllable is less than the minimum syllable duration, the onset and offset pair is removed.
6. If any onsets or offsets are outside of the time-range of interest (range determined by the slider below the binary image), they will be removed.
7. If the segmentation needs a small adjustment, such as a missing onset or offset or an incorrect placement due to noise, the user can manually add or delete onsets and offsets.
8. Lastly, the user can submit the parameters, the final binary matrix, and syllable onsets and offsets. If a satisfactory segmentation was not reached, the file can be tossed.

#### Quantitative Analysis

Once syllable segmentation has been completed in Chipper, output files (gzips) are generated containing all necessary information on the binary image, segmentation, and conversion factors for both time and frequency space. These output files can then be processed using Chipper’s analysis tool. This portion of Chipper is fully automated; the window serves to show the number of files processed out of the total selected by the user. For each song being processed, Chipper produces multiple song, syllable, note, and syntax measurements (**Table 1**). Many of these outputs rely on the input parameters for noise and syllable similarity thresholds; thus, we recommend using our widgets in Chipper to determine appropriate thresholds for each species-specific set of songs studied. When the quantitative song analysis is complete, the user can perform statistical tests on the quantitative song measures for a particular research question of interest.

**Table 1.** Chipper’s output measurements.

| Measurement | Calculation |
| --- | --- |
| Song duration | (time of last syllable offset – time of first syllable onset) |
| Number of syllables | number of syllable onsets in a song |
| Syllable rate | (number of syllables)/(song duration) |
| Average syllable duration | mean(time of syllable offset – time of syllable onset) |
| Standard deviation of syllable duration | standard deviation(time of syllable offset – time of syllable onset) |
| Average silence duration | mean(time of syllable onset – time of previous syllable offset) |
| Standard deviation of silence duration | standard deviation(time of syllable onset – time of previous syllable offset) |
| Largest syllable duration | max(time of syllable offset – time of syllable onset) |
| Smallest syllable duration | min(time of syllable offset – time of syllable onset) |
| Largest silence duration | max(time of syllable onset – time of previous syllable offset) |
| Smallest silence duration | min(time of syllable onset – time of previous syllable offset) |
| Average syllable frequency modulation | mean(maximum frequency – minimum frequency for each syllable) |
| Std. dev. syllable frequency modulation | standard deviation(maximum frequency – minimum frequency for each syllable) |
| Average syllable lower frequency | mean(minimum frequency of each syllable) |
| Average syllable upper frequency | mean(maximum frequency of each syllable) |
| Largest syllable frequency modulation | max(maximum frequency – minimum frequency for each syllable) |
| Smallest syllable frequency modulation | min(maximum frequency – minimum frequency for each syllable) |
| Maximum syllable frequency | max(maximum frequency of each syllable) |
| Minimum syllable frequency | min(minimum frequency of each syllable) |
| Overall syllable frequency range | max(maximum frequency of each syllable) – min(minimum frequency of each syllable) |
| Syllable stereotypy | list of the mean(pairwise percent similarities) for each repeated syllable, where percent similarity is the maximum(cross-correlation between each pair of syllables)/maximum(autocorrelation of each of the compared syllables) × 100 |
| Mean syllable stereotypy | mean(stereotypy values for each repeated syllable) |
| Standard deviation syllable stereotypy | standard deviation(stereotypy values for each repeated syllable) |
| Number of unique syllable | number of unique values in syllable pattern |
| Degree of repetition | (number of syllable onsets in a song)/(unique values in syllable pattern) |
| Syllable pattern | list of the syllables in the order that they are sung, where each unique syllable is assigned a number (i.e. the song syntax) |
| Degree of sequential repetition | number of syllables that are followed by the same syllable/(number of syllables - 1) |
| Number of notes | number of 4-connected elements of the spectrogram with an area greater than the noise threshold |
| Number of notes per syllable | (total number of notes)/(total number of syllables) |
| Average note duration | mean(time of note beginning – time of note ending) |
| Standard deviation of note duration | standard deviation(time of note beginning – time of note ending) |
| Largest note duration | max(time of note beginning – time of note ending) |
| Smallest note duration | min(time of note beginning – time of note ending) |
| Overall note frequency range | max(maximum frequency of each note) – min(minimum frequency of each note) |
| Average note frequency modulation | mean(maximum frequency – minimum frequency for each note) |
| Std. dev. note frequency modulation | standard deviation(maximum frequency – minimum frequency for each note) |
| Average note lower frequency | mean(minimum frequency of each note) |
| Average note upper frequency | mean(maximum frequency of each note) |
| Largest note frequency modulation | max(maximum frequency – minimum frequency for each note) |
| Smallest note frequency modulation | min(maximum frequency – minimum frequency for each note) |
| Maximum note frequency | max(maximum frequency of each note) |
| Minimum note frequency | min(minimum frequency of each note) |

### Additional note and syntax analyses for birdsong application

For a subset of measurements provided by Chipper’s analysis tool, the user can improve measurement accuracy by setting a noise threshold and syllable similarity threshold. The noise threshold affects any note-related and frequency-related calculations, since any signal smaller than the noise threshold is removed from the binary spectrogram. For example, low-frequency noise in a syllable that is not removed either in the segmentation process or by the noise threshold will affect multiple frequency measurements (minimum syllable frequency, average syllable frequency modulation, etc.) Any calculations that specifically uses the onsets and offsets, such as song, syllable, and silence durations, will not be affected by the noise threshold. The syllable similarity threshold only affects syntax-related calculations (number of unique syllables, syllable pattern, syllable stereotypy, etc.). As it is useful to set these thresholds based on multiple songs the user is processing, we have provided widgets to visualize the choice of these thresholds, described below.

#### Determine threshold for noise

Chipper’s quantitative analysis uses connectivity to classify signal within a syllable as either a note or noise. Specifically, any signal within the syllables (defined by onsets and offsets) in the binary image that is connected by an edge (not corner, i.e. 4-connected) and has an area greater than the user-specified threshold is labeled as an individual note, and any signal with an area less than or equal to the threshold is considered noise. Since this division in signal is highly dependent on signal-to-noise ratio or amplitude, we provide a widget to determine the best threshold for a set of songs. In the noise threshold widget, the user selects a folder of multiple gzips (the output from syllable segmentation) for a representative subset of the songs being analyzed. For each song, the user selects a noise threshold. The user can change the threshold to visualize areas being classified as notes (colored) versus noise (white) within the song, and then submits a threshold for that song. After going through the selected songs, a summary is provided including the average, minimum, and maximum thresholds selected for noise and, if enough songs are processed in the widget, a histogram. This information is provided to guide the user in choosing a single threshold that will be used in song analysis for the entire set of song files. We advise caution in using the output from note analysis for low-quality recordings: whereas high-quality recordings will have syllables in which signal is only disconnected at true notes, the degraded signal in low-quality recordings can lead to many false notes.

#### Determine threshold for syllable similarity

For each pair of syllables, a percent syllable similarity is calculated by sliding one syllable’s binary matrix across another syllable’s binary matrix and finding the maximum overlap (cross-correlation). This is then repeated for each syllable compared to itself, providing an autocorrelation for each syllable. We scale the maximum overlap between the two syllables by dividing by the maximum of the two syllables’ autocorrelations; multiplied by 100, this results in a percent of the maximum possible overlap or percent syllable similarity for the syllable pair. Similar methods of spectrographic cross-correlation have been previously demonstrated as a useful method in determining syllable types (Clark, Marler & Beeman 1987). Applying the user-defined syllable similarity threshold to the resulting pairwise matrix, we establish the syntax for the recording by considering two syllables the same if their similarity is greater than or equal to the user-specified threshold. If two syllables are considered to be the same type and the second one of those is considered the same as a third syllable, then the third syllable is classified as the same type as the first two. This prevents groups of similar syllables from being separated but also means that the first and third syllables could have a percent similarity below the threshold but still be considered the same type. Chipper’s syllable similarity threshold widget guides the user in deciding an appropriate value. The binary song and the corresponding syllable onset and offset lines from syllable segmentation are plotted. Based on the threshold, the syntax is displayed in text as well as visually, with like syllables colored the same. The user can change the threshold to see how this will change the syntax of the song. When all of the sample songs have been processed, a summary will be displayed with the average, minimum, and maximum thresholds selected for syllable similarity and, if enough songs have been processed, a histogram. Once again, this information is provided to guide the user in choosing a threshold to process the entire set of song files of interest.

## Conclusions

With the ever-growing repositories of citizen-science recordings, a new software developed specifically to handle the varying recording qualities and vast species coverage was needed (**Figure 3**). Thus, we developed Chipper as a free, open-source software to improve the workflow of audio signal processing with particular application to high-throughput analysis of citizen-science recordings. With its user-friendly graphical user interface, Chipper can be used by researchers, students in classrooms, and curious citizen-scientists alike. In testing Chipper, we found that it produced robust estimates of sound properties for a set of synthetic recordings, and these results were consistent within and between users and in the presence of natural and white noise (**Supporting Information**). We hope Chipper, in tandem with citizen-science data, can aid in large-scale spatiotemporal studies of acoustic signals, particularly global inter- and intra-species studies of birdsong. Additionally, since Chipper has open-source code on GitHub, users can extend and contribute to Chipper, tailoring it to additional projects and data types. Ultimately, using Python and Kivy, we have developed an application that facilitates audio processing of natural recordings, extending the utility of rapidly growing citizen-science databases and improving the workflow for current birdsong research in ecology and evolution.

**Figure 3.**
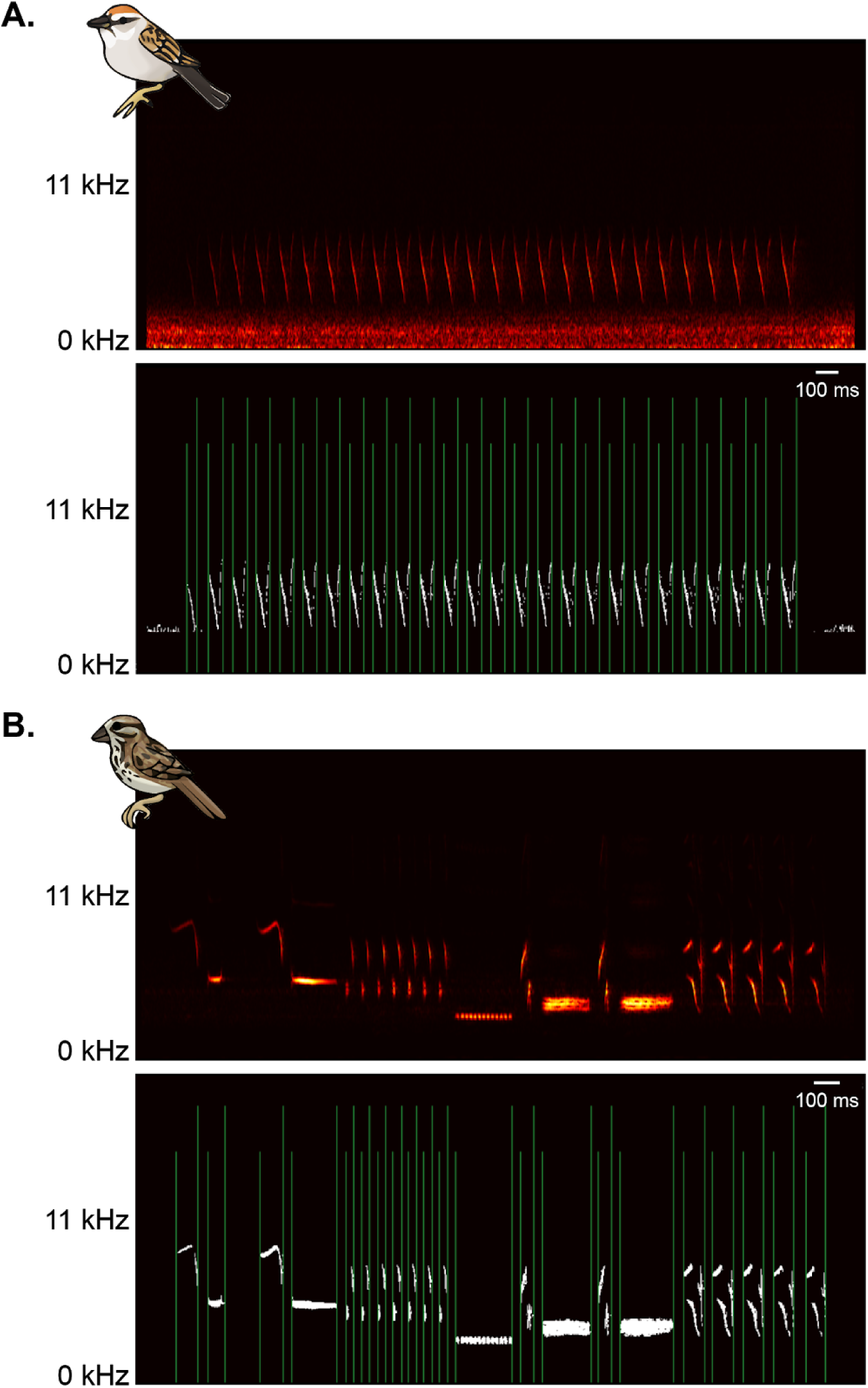
Chipper can segment songs of various qualities and from different species. Example song of (A) chipping sparrow and (B) song sparrow. The top images are the spectrograms when they are originally loaded into Chipper. The bottom images are the binary signal after parameters have been adjusted to optimize segmentation. The green lines show the onsets (shorter lines) and offsets (longer lines) for each syllable.

## Supporting information

Chipper Manual

## Author Contributions

A.M.S and N.C. conceived and designed the project. A.M.S. and J.C.P. developed the software. Parts of the software were based on earlier code written by N.C. in MATLAB; A.M.S. translated this code to Python and built the graphical user interface in Kivy. J.C.P. packaged the software. A.M.S. led the writing of the manuscript with assistance and revision from J.C.P. and N.C. All authors contributed to drafts, edited the final manuscript, tested and verified the software, and gave final approval for publication.

## Acknowledgements

We thank the following students for testing Chipper during its development: Vanderbilt University BSCI 1512L Fall 2018 class, Vanderbilt University BSCI 3965 Spring 2019 class, Megan Mitchell, Nyssa Kantorek, Maria Sellers, and Emily Beach. In addition, we thank Megan Mitchell for editing the Chipper Manual and Cristina Robinson for the bird art used in this paper and in the Chipper logo.

## Data Accessibility

Chipper and code to create and analyze synthetic songs can be found at https://github.com/CreanzaLab/Chipper.

The recordings used in this paper are freely available in the Xeno-Canto repository: Jonathon Jongsma, XC320440. Accessible at www.xeno-canto.org/320440; Chris Parrish, XC13690. Accessible at www.xeno-canto.org/13690; Allen T. Chartier, XC16985. Accessible at www.xeno-canto.org/16985.

The folder and file images used in Figure 1 are adapted from icons found at the Noun Project: tab file document icon by IYIKON, .WAV Folder by Linseed Studio, Audio by Ben Avery, zip file document icon by IYIKON, wax file document icon by IYIKON, and csv file document icon by IYIKON.

## Software availability

Chipper v1.0 can be downloaded for Mac, PC, and Linux at https://github.com/CreanzaLab/Chipper/releases. Chipper leverages several existing Python packages including SciPy (Jones, Oliphant & Peterson 2001), Pandas (McKinney 2010), Matplotlib (Hunter 2007), and NumPy (Oliphant 2006; van der Walt, Colbert & Varoquaux 2011). We also use the Python library Kivy v.1.10 for building the graphical user interface (Virbel, Hansen & Lobunets 2011).

