## Supporting Information for "Chipper: Open-source software for semi-automated segmentation and analysis of birdsong and other natural sounds"

*Generation of a standardized test set of synthetic birdsongs*

We used the SciPy module in Python to generate 50 unique synthetic birdsongs (function chirp). Each song has 10 syllables in the following order: linearly constant, linearly increasing, linearly decreasing, quadratically increasing and concave, quadratically increasing and convex, quadratically decreasing and convex, quadratically increasing and concave, symmetric quadratic, logarithmically increasing, and logarithmically decreasing. For each song, an amplitude was randomly chosen from a uniform distribution between 100–10,000. Then, within a song, an amplitude for each syllable was randomly selected from a uniform distribution of amplitudes 30% above or below the song amplitude. The amplitude of each syllable was then altered to linearly increase to the maximum amplitude over the first 40% of the syllable length and then linearly decrease to zero over the last 40% of the syllable length; this smoothing both mimics natural signals and avoids discontinuities when performing the fast fourier transform on the waveform to produce the spectrogram. Lastly, the amplitude of each syllable was multiplied by an exponential decay function to mimic the natural decrease in signal intensity in bird sounds. Similarly, the starting frequency was randomly selected from a discrete, uniform distribution between 2,000–10,000 Hz; the ending frequency was then either the same as the starting frequency for flat syllables or randomly selected from the range of 2,000 Hz to starting frequency or from the range of the starting frequency to 10,000 Hz, depending on the shape of the syllable. The syllable lengths and the lengths of silence between each syllable were also randomly selected from 0.1–0.9 seconds and 0.01–0.5 seconds, respectively. Lastly, the beginning and ending of each generated song was padded with ~200ms of silence. Each song was saved as a WAV file, and all corresponding syllable and silence parameters were saved in a text file.

Next, different types of noise were added to each of the 50 synthetic songs. Using Audacity (Generate > Noise), two different tracks of white noise were created with 0.01 and 0.001 amplitude. Two tracks of natural noise were created by selecting sections of two different passive recordings collected with a Wildlife Acoustics Song Meter 4 recorder that had minimal extraneous sounds (e.g. birds, crickets, car horns, etc.). Each of these noise tracks (white or natural) were added to the synthetic songs, creating 4 different noisy recording sets of 50 songs each (**Supplemental Figure 1**). The maximum amplitude of the noise and the signal for each song was calculated.

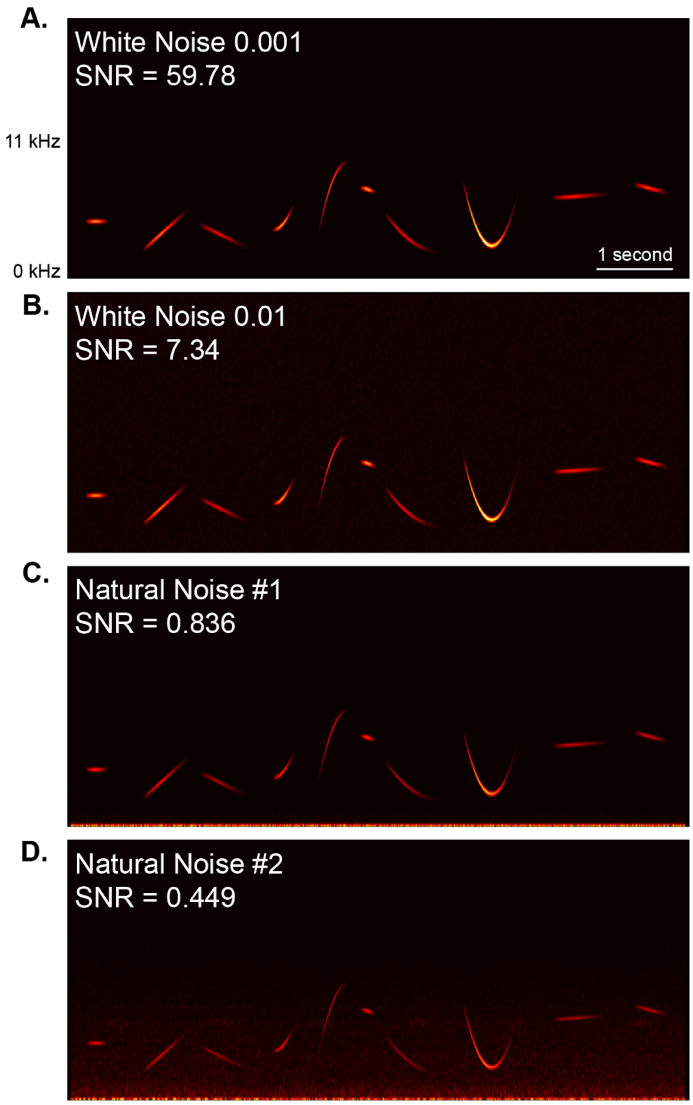


**Supplemental Figure 1. Example of a synthetic song with the four levels of added noise.** (A) White noise with a very low amplitude of 0.001 (in Audacity) was added to the synthetic song. It is visually apparent that the signal-to-noise ratio (SNR) for this song is very high at 59.78. (B) A white noise track with a higher amplitude 0.01 (in Audacity) was added to the synthetic song, lowering the SNR for this song to 7.34. (C) The Natural Noise #1 track has most of the signal in the low-frequency range. As the noise is very close in amplitude to the signal, the SNR drops to 0.836; however with Chipper the high-pass filter can remove most of this noise easily. (D) Similarly Natural Noise #2 (SNR 0.449) has a significant amount of high amplitude noise in the low-frequency range. The high-pass filter can easily remove this band of noise, but there is still significant noise at higher frequencies occupied by the song.

*Assessing accuracy, repeatability, and reproducibility of Chipper with synthetic birdsongs*

All 250 synthetic songs (1 set without noise, 2 sets with natural noise, and 2 sets with white noise) were then processed in Chipper independently by two users. First, we compared the measurements from Chipper for the set of synthetic songs without added noise to the actual values used to create the songs (**Supplemental Figure 2A and 3A**). Since we created the syllables with amplitude linearly increasing over the first 40% of the syllable and decreasing over the last 40% to mimic real birdsongs, it was not surprising that the measured syllable durations are shorter than the “actual” values since the syllables begin and end at very low amplitude (**Supplemental Figure 2A, row 2**). Similarly, the opposite effect occurs for average silence durations (**Supplemental Figure 3A, row 1**). Our baseline measurements for bout duration and average syllable frequency modulation are close to the actual values used to generate the synthetic songs (**Supplemental Figure 2A, row 1 and 3**). Next, we compared the measurements from Chipper for the sets with noise to the measurements from Chipper for the set without noise. Specifically, we calculated a signal to noise ratio for each song using the maximum amplitude of the noise and the maximum amplitude of the signal that was documented when creating the synthetic songs. Then, we subtracted the measurements from Chipper for the noisy files from the measurements from Chipper for the set of songs without noise, which we used as the baseline measurements (**Supplemental Figure 2B and 3B**). For most measurements we assessed, the discrepancy between noisy files and files without noise began to increase when signal amplitude was approximately less than twice the amplitude of noise. Lastly, we tested the repeatability and reproducibility of Chipper measurements. The measurements from Chipper for all sets of synthetic songs were compared between users (**Supplemental Figure 2C and 3C**). Then the measurements from Chipper for all sets of synthetic songs were compared between the same user’s first and second attempts at segmenting the songs in Chipper (**Supplemental Figure 2D and 3D**).

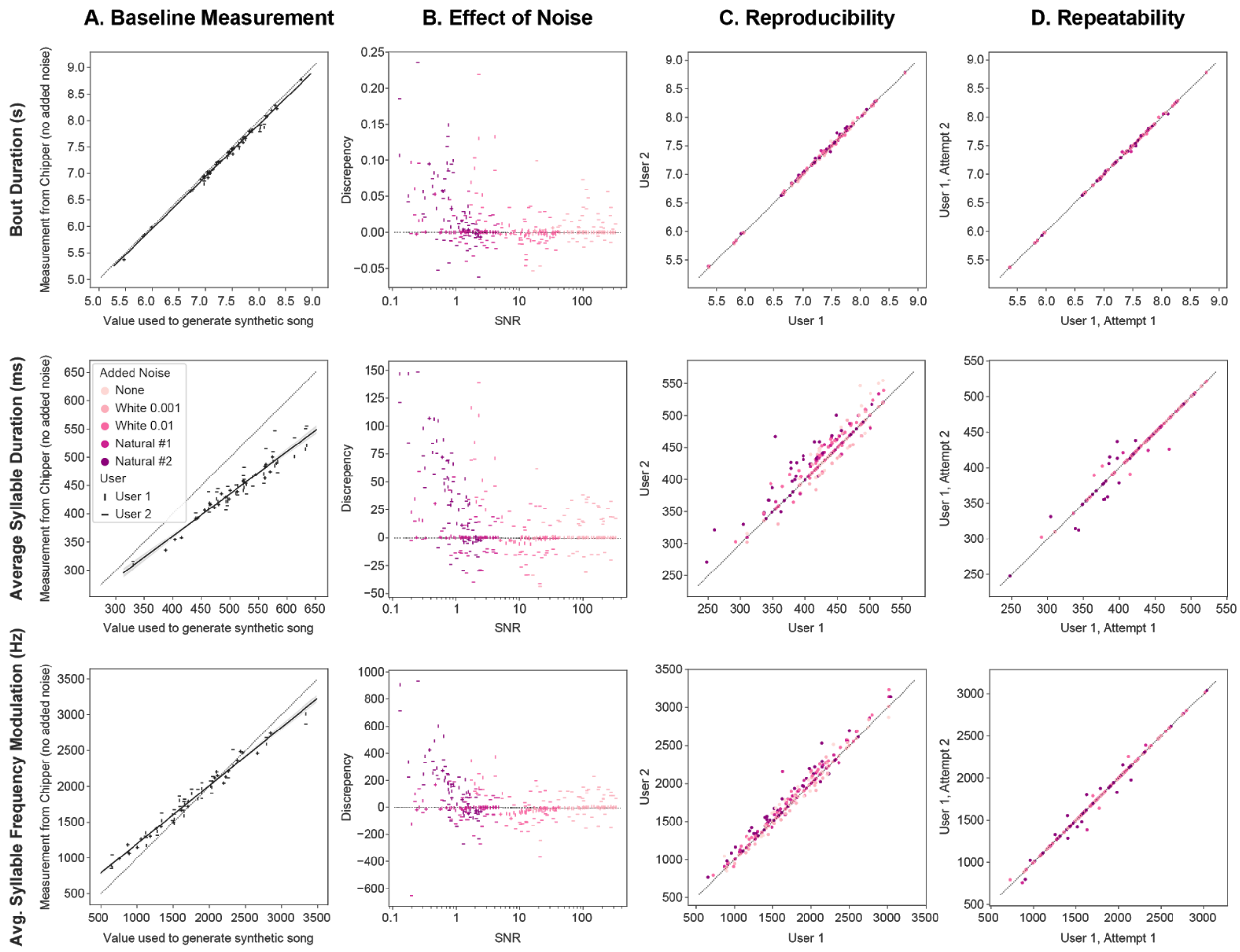


**Supplemental Figure 2. Accuracy, repeatability, and reproducibility of Chipper for key song features.** (A) We compare the measurements from Chipper segmentation and subsequent analysis (y-axis) to the actual values (x-axis) used to generate the synthetic songs with no added noise. This provides a baseline for accuracy, as we expect measurements will not align with the line of unity (shown as dotted line) due to the method of creating synthetic songs. (B) We examine the effect of both white noise and natural noise on the segmentation and analysis of synthetic songs in Chipper. The discrepancy (y-axis) is the difference between the Chipper-measured song feature from synthetic songs with added noise and from songs without noise. The signal-to-noise ratio (SNR) can be indicative of the possible accuracy; however, natural noise often has low-frequency noise that Chipper can reduce, providing a better accuracy than expected solely based on SNR. (C) To assess reproducibility, the measurements from segmentation and subsequent analysis conducted by two users is compared for synthetic songs of various noise levels. (D) To assess repeatability, the measurements from segmentation and subsequent analysis conducted twice by the same user are compared for all synthetic songs. These assessments are shown for three song features (rows): bout duration, average syllable duration, and average syllable frequency modulation. Similar plots for additional song features can be found in **Supplemental Figure 3**.

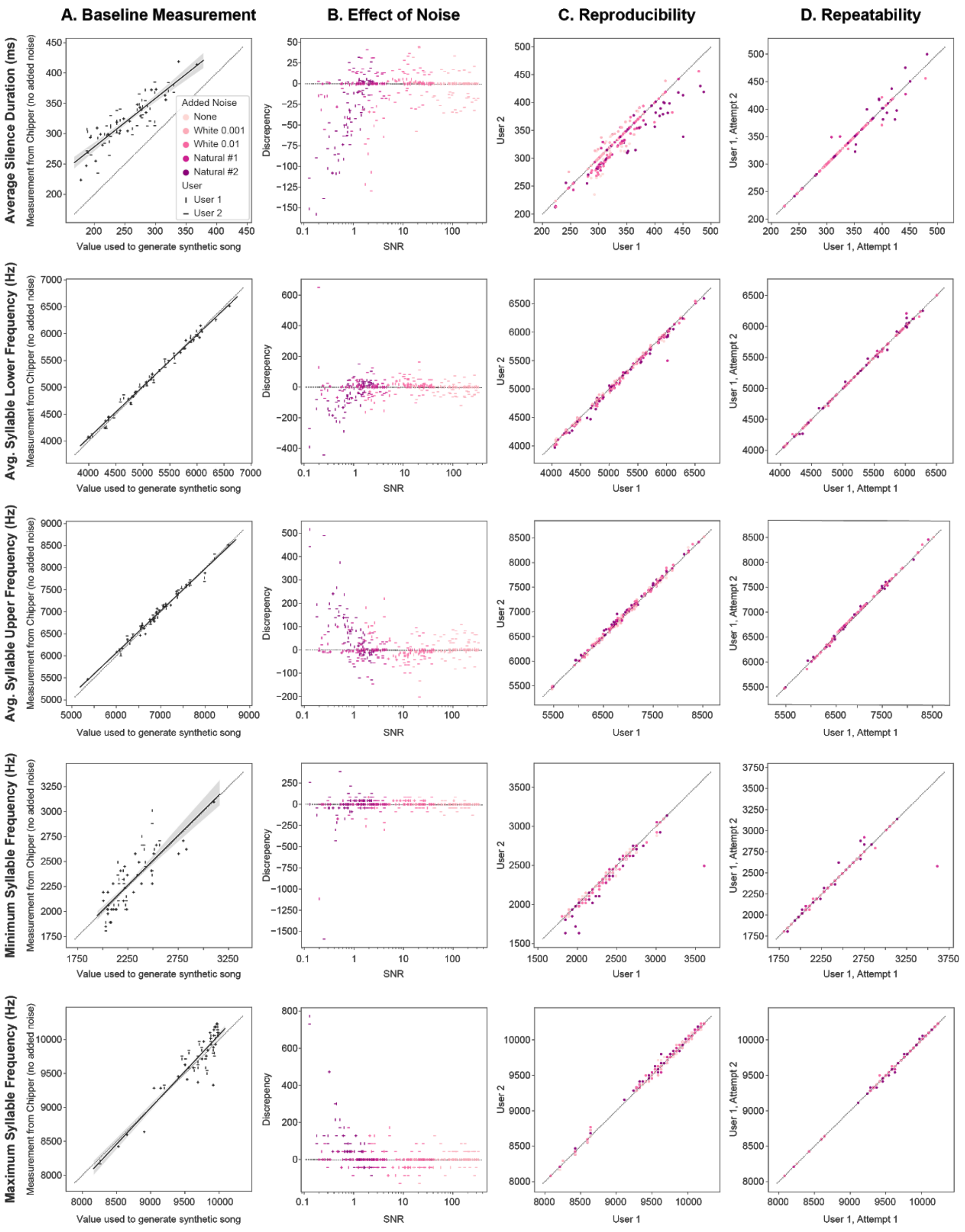


**Supplemental Figure 3. Accuracy, repeatability, and reproducibility of Chipper for additional song features.** (A-D) See Supplemental Figure 2 for description of plots. Assessments are shown for five song features (rows): average silence duration, average syllable lower frequency, average syllable upper frequency, minimum syllable frequency, and maximum syllable frequency.
