## Supplementary material for "Chipper: Open-source software for semi-automated segmentation and analysis of birdsong and other natural sounds": Chipper Manual

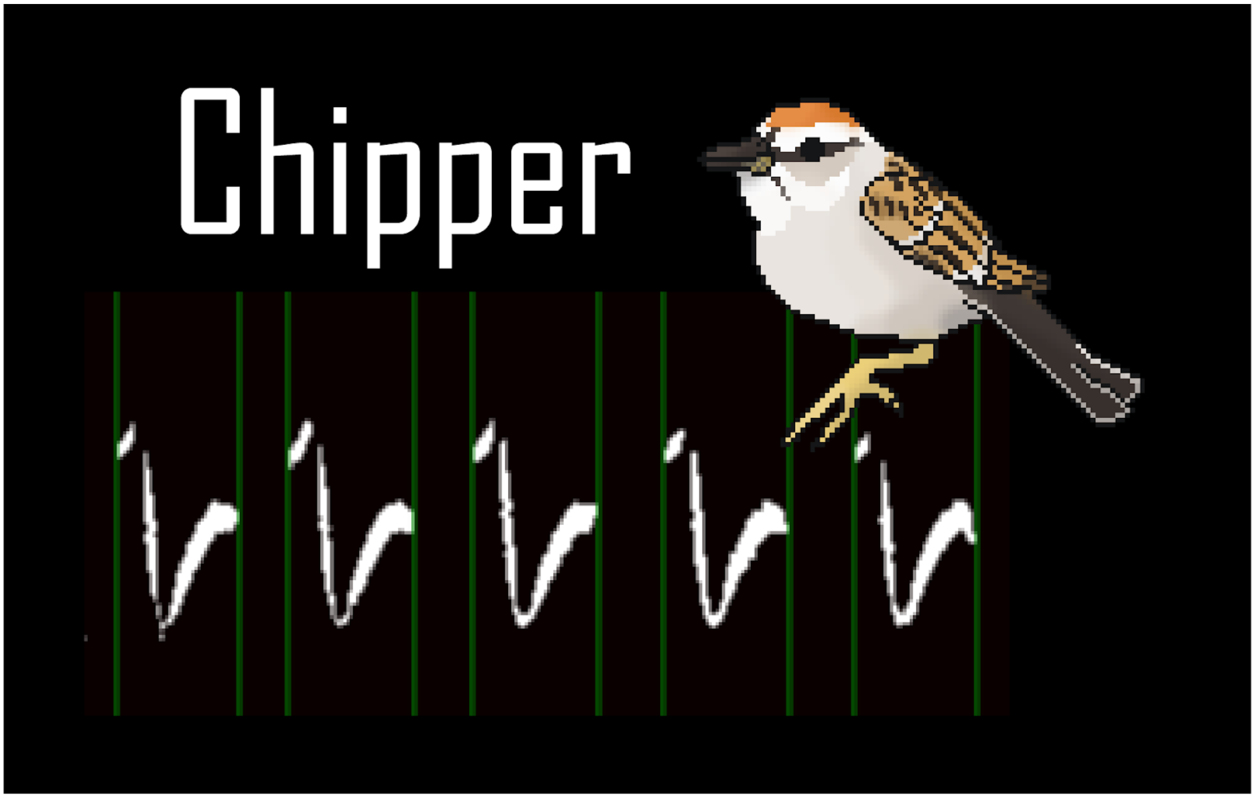
Chipper

**Table of Contents**


1. How to Install
2. Overview
3. Syllable Segmentation
4. Example of Syllable Segmentation
5. Noise Threshold Widget
6. Syllable Similarity Threshold Widget
7. Running Analysis
8. Analysis Output

**How to Install**

Option 1: Download Chipper

1. Download the version of Chipper for your operating system (found at https://github.com/CreanzaLab/chipper/releases).
2. Unzip the folder.
3. Navigate to and double click the executable/application named run_chipper, which may have a bird as an icon.
4. You will now see a terminal window open, and soon after, the Chipper landing page will load. You are ready to go!

Option 2: Install from source (primarily for developers)

1. Install Anaconda
   Our recommended approach is to use Anaconda, which is a distribution of Python containing most of the numeric and scientific software needed to get started. If you are a Mac or Linux user, have used Python before and are comfortable using pip to install software, you may want to skip this step and use your existing Python installation.

2. From within a terminal or command prompt window we will install most packages with conda

```
$ conda create -n chipper_env python=3.7
$ condapip install  pypiwin32 kivy.deps.sdl2 kivy.deps.glew
$ kivy.deps.gstreamer kivy.deps.glew_dev kivy.deps.sdl2_dev
$ kivy.deps.gstreamer_dev
```

3. Additional steps for Windows users:

```
$ pip install  pypiwin32 kivy.deps.sdl2 kivy.deps.glew
kivy.deps.gstreamer kivy.deps.glew_dev kivy.deps.sdl2_dev
kivy.deps.gstreamer_dev
```

4. Install kivy packages

```
$ garden install --kivy graph
$ garden install --kivy filebrowser
$ garden install --kivy matplotlib
$ garden install --kivy progressspinner
```

For full code visit https://github.com/CreanzaLab/chipper.

**Overview**

The Chipper landing page allows you to choose whether you want to

1) segment songs (or other acoustic signals)
2) determine the threshold for noise
3) determine the threshold for syllable similarity, or
4) complete analysis of already segmented files.

These steps should generally be conducted in the order listed. Steps 2 and 3 can be skipped if you have already determined appropriate thresholds for your study.

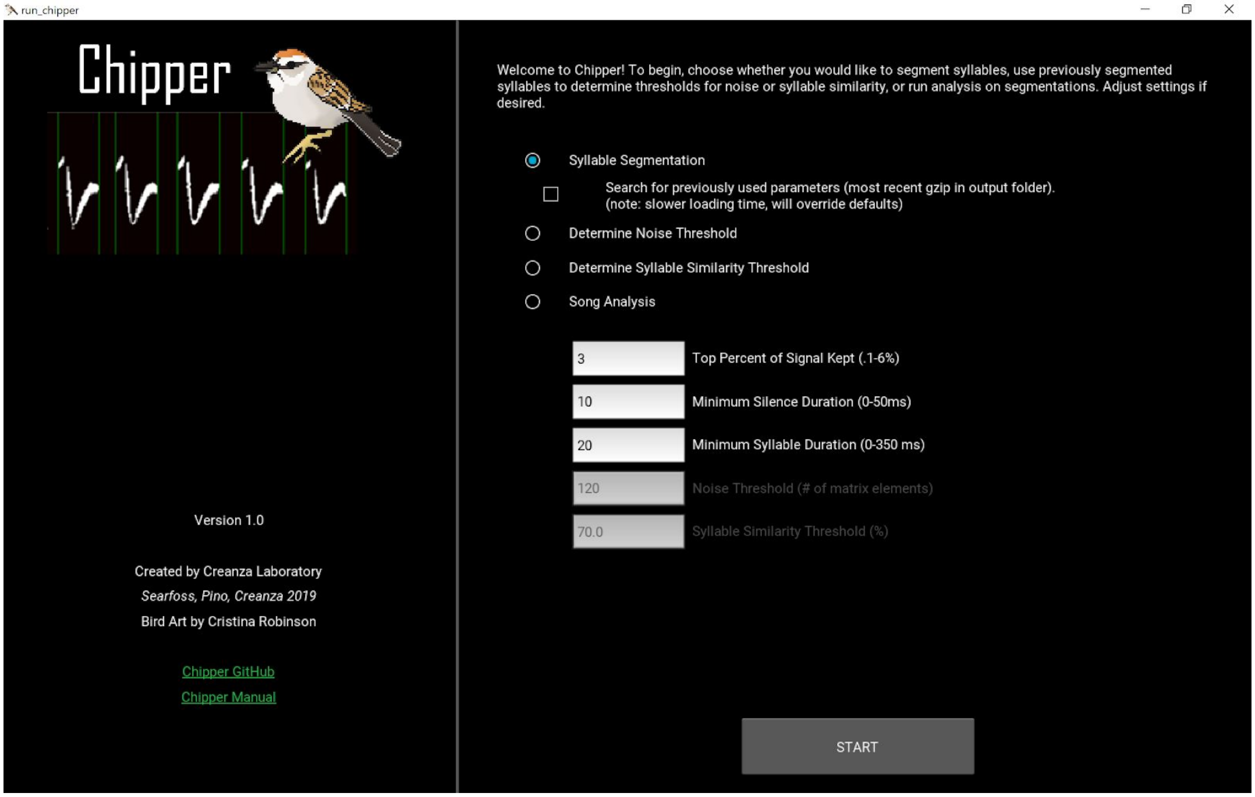


For **Syllable Segmentation**, the user can choose parameter defaults for top percent of signal kept, minimum silence duration, and minimum syllable duration; these defaults will start the parsing of every song. You can also select “Search for previously used parameters” which will look for gzips in any *SegSyllsOuput_YYYYMMDD_THHMMSS* folders in the selected folder for segmentation. If any gzips are found, the most recent one will be used to load the previous settings and segmentation conducted for the song.

For **Song Analysis**, the user can choose parameters for noise threshold (to distinguish between signal and noise) and percent syllable similarity (to determine syntax). We suggest first using the noise and syllable similarity threshold widgets for best results; after finishing each widget, the new parameter for analysis will be populated.

**Syllable Segmentation**

Starting Syllable Segmentation will first take you to a file explorer to choose either a single WAV file or folder of WAV files. The length of the songs and the size of the files Chipper can handle depends greatly on your computing resources and screen size. We recommend songs between ~0.5s and 10s long or files no larger than 3MB. Depending on the computing resources, you may experience lag with files between 2-3MB. Thus, we recommend choosing bouts of song to segment using another program such as Audacity (audacity.sourceforge.net). Similarly, if a bout is very short, it may appear stretched; you can always reduce this effect when selecting a bout in Audacity by not cutting the bout too close or by adding time to the beginning or end of the bout.

*Note: We have set a warning message for files over 3MB in which the user can select to either toss or process the file; this is a safety to ensure the user knows they may experience a lag or even crash Chipper if the file is much larger than recommended and has not been previously parsed. If you are consistently parsing files larger than this with no issue and want to change this threshold, see line 338 of control_panel.py (and line 275 in run_chipper.kv for popup message).*

Each file will load using the default parameters from the landing page to automatically parse the song. Next adjust the sliders to finalize your segmentation. For detailed calculations involving the parameters underlying these sliders and how they are applied to the spectrogram, see *Searfoss, Pino, Creanza 2019* or the code at github.com/CreanzaLab/chipper.

Here is a suggested order in which to adjust the sliders (for a detailed example with screenshots, see next section):

1. Adjust the high-pass and low-pass filter slider (left of top spectrogram).

2. If there are some portions of the song that seem very low in amplitude, try normalizing the signal.

3. Adjust the signal threshold to reduce noise.

   *Tip: You can use the crop sliders (under the bottom spectrogram) to remove the onsets/offsets temporarily to see if you like what the signal threshold is giving you without having the lines crowd your view. Be sure to put these sliders back after you are done, as the onsets and offsets must be in place to submit a segmentation.*

4. Adjust minimum silence duration to the ***minimum value*** that segments the way you think is correct. Using the smallest value that gives you good parsing will help if you have spots of noise right on the edge of a syllable; it will cut the speck of noise out of the actual syllable. On the other hand, if you are trying to get two parts of a syllable that are

far apart from one another to parse together, try increasing the minimum silence duration quite a bit.

5. You may need to iterate between steps #3 and #4.

6. Adjust the minimum syllable duration if you are still not satisfied with the parsing. Usually you will not have to adjust this parameter much or at all. (However, this will depend on the song type you are parsing.) It is often useful in getting rid of little bits of noise that are parsing as syllables (by increasing the minimum syllable duration) or to include small syllables that are not parsing (by decreasing the minimum syllable duration).

7. Use the crop sliders (under the bottom spectrogram) to remove onsets and offsets capturing any signal before or after the song.

8. Add/Delete any onsets/offsets that are missing or extraneous. Try to add as few manual onsets/offsets as possible. It is better to have too many and have to remove.

   *Tip: Use the left and right arrow keys to move between the selected onset/offset and the others. Use "Enter" to accept addition or deletion of onsets/offsets. Use "x" to cancel addition or deletion of onsets/offsets.*

9. Submit your parsed song, or toss if you think it is too noisy and you are not getting good data!

   *Careful not to hit submit or toss twice in a row. If you think the button did not work, it might just be loading the next file. The **buttons should turn blue if they have been selected** and will be gray again when the new image is loaded. However, if you do accidentally skip or toss a song, you can use the "Back" button to return to the previous song.*

Once you have parsed all the files in a folder, there will be a new folder within that directory called *SegSyllsOutput_YYYYMMDD_THHMMSS*, where *YYYYMMDD* is replaced with the current date and *HHMMSS* is replaced by the current time. For every WAV file that was successfully segmented, there will be an associated gzip which will be used in the next steps in Chipper. Specifically, each gzip is written after the user hits "Submit". These gzips are used to review the segmentation again using Syllable Segmentation or to determine Thresholds and run Analysis in the next steps.

In addition, four human-readable text files are output once the last file is either submitted or tossed.

1. *segmentedSyllables_parameters_all* includes a list of all the Chipper parameters used to reach the submitted segmentation.

| | |
|---|---|
| FileName | name of WAV file |
| FrequencyFilter | [high-pass, low-pass] in number of matrix elements from the bottom of the spectrogram |
| BoutRange | [left crop, right crop] in number of matrix elements from the left of the spectrogram |
| PercentSignalKept | top percent of signal kept |
| MinSilenceDuration | number of spectrogram matrix elements |
| MinSyllableDuration | number of spectrogram matrix elements |
| Normalized | 'yes' or 'no' indicating whether the song was normalized or not |

2. *segmentedSyllables_syllables_all* with a list of onsets and a list of offsets in number of matrix elements from the left of the spectrogram.

3. *segmentedSyllables_conversions_all* includes the conversions necessary to change the parameters from spectrogram matrix elements into milliseconds or Hz for each WAV file processed.

| | |
|---|---|
| FileName | name of WAV file |
| timeAxisConversion | number of milliseconds per matrix element |
| freqAxisConversion | number of Hz per matrix element |

4. *segmentedSyllables_tossed* with a list of the files that were tossed. There should be no gzips provided for the files in this list.

*Note: If Chipper crashes during segmentation (or the user exits Chipper), the text files will not be output; however, gzips for any segmentations that were submitted will be present in the output folder. To run the next steps, only gzips are needed. The text outputs are solely for the user's information.*

### Example of Syllable Segmentation

1. Select “Syllable Segmentation”.

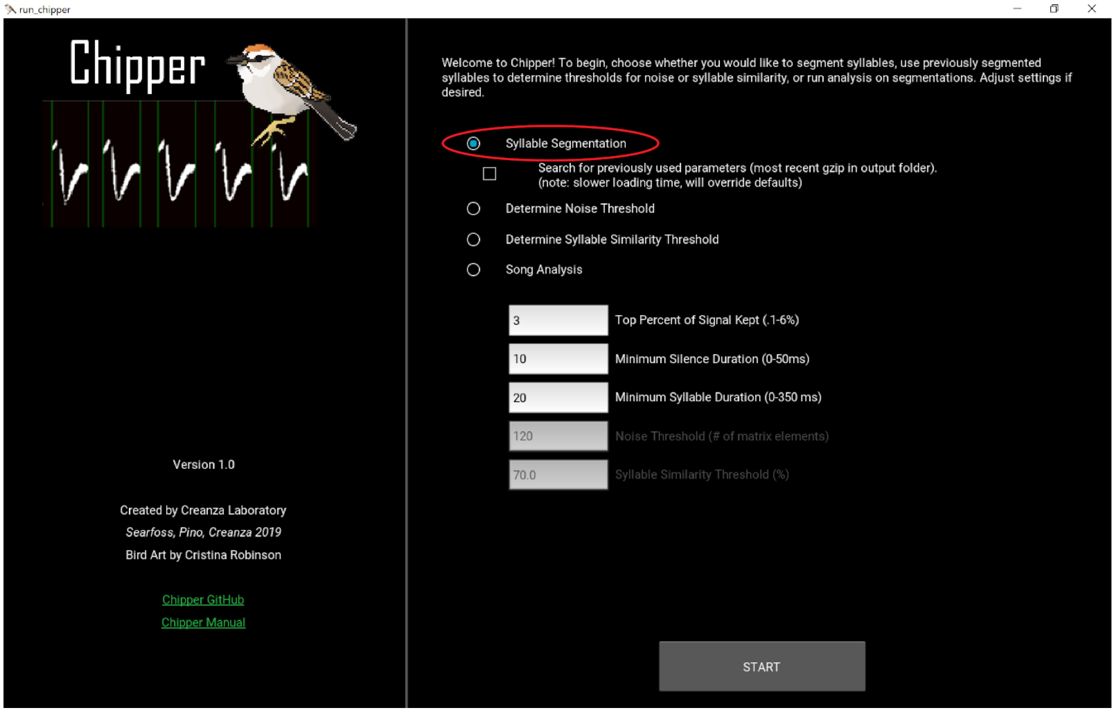


2. Navigate to and select a WAV file or folder of WAV files to parse. Double click ..\ (for PC) or ../ (for Mac or Linux) to go back up a folder. Here we use the second bout from the Xeno Canto recording 381923. (XC381923 contributed by Lucas, Creative Commons Attribution-NonCommercial-ShareAlike 4.0, https://www.xeno-canto.org/381923).

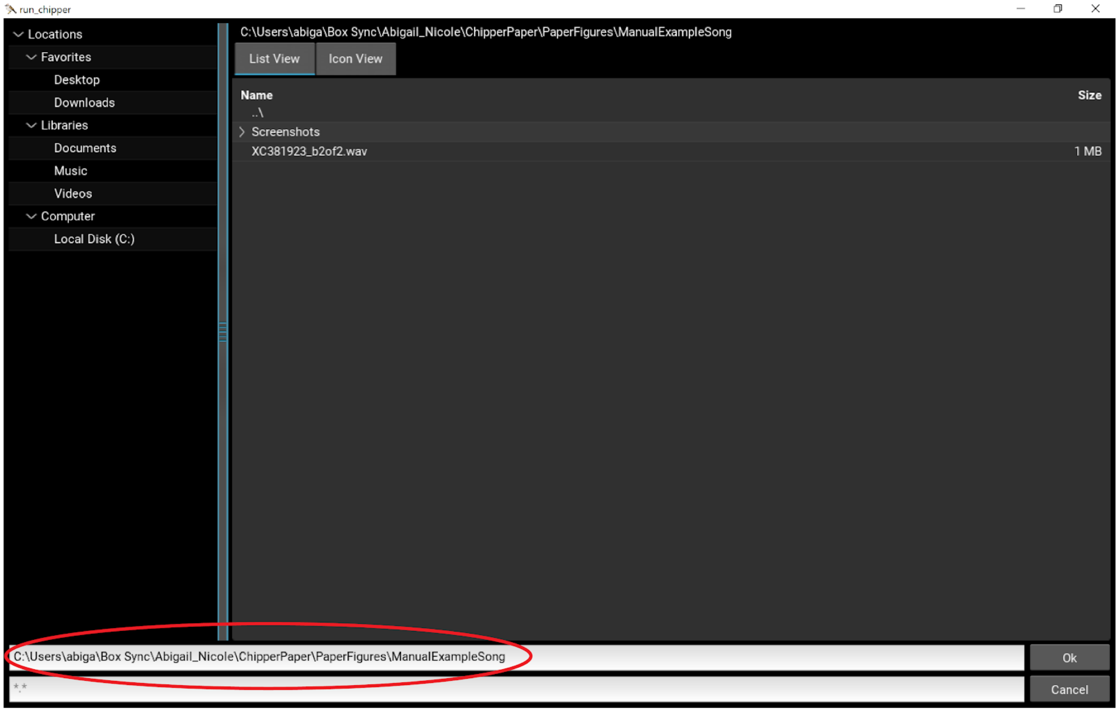

3. Click “Begin Segmentation”.

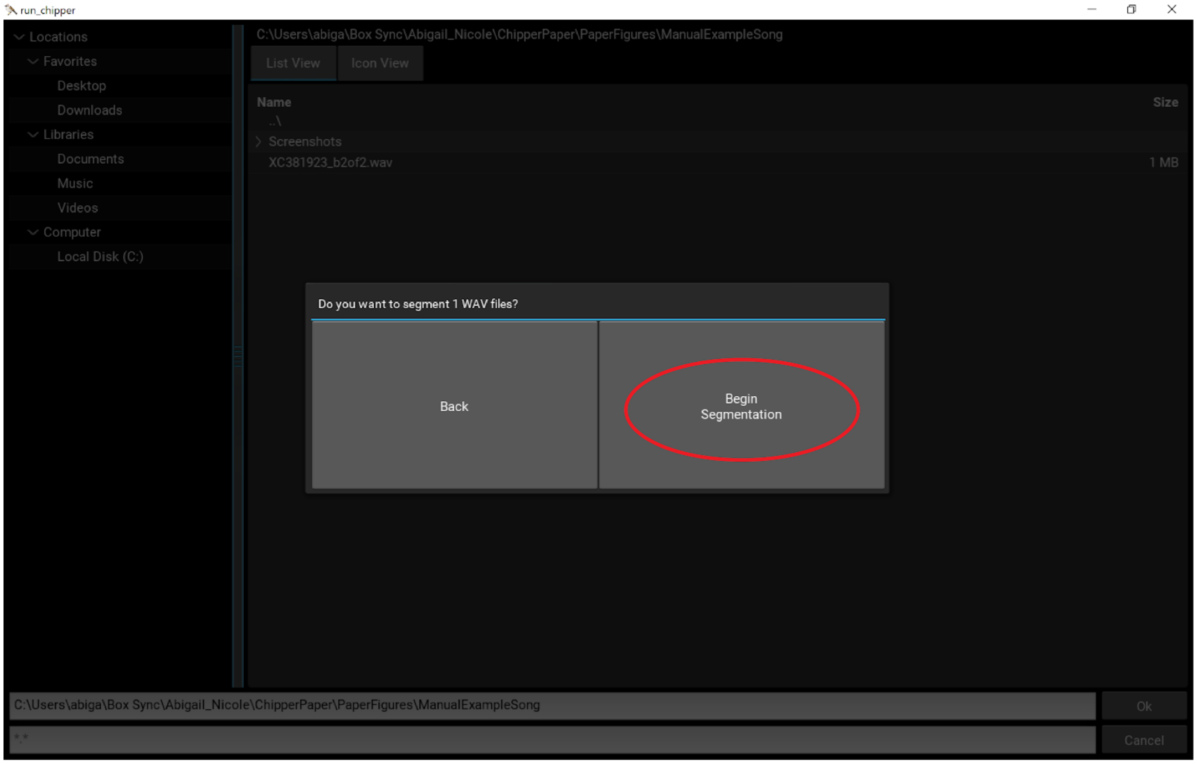


4. The file is loaded with segmentation using the default parameters. In this case, we did not change the defaults on the landing page.

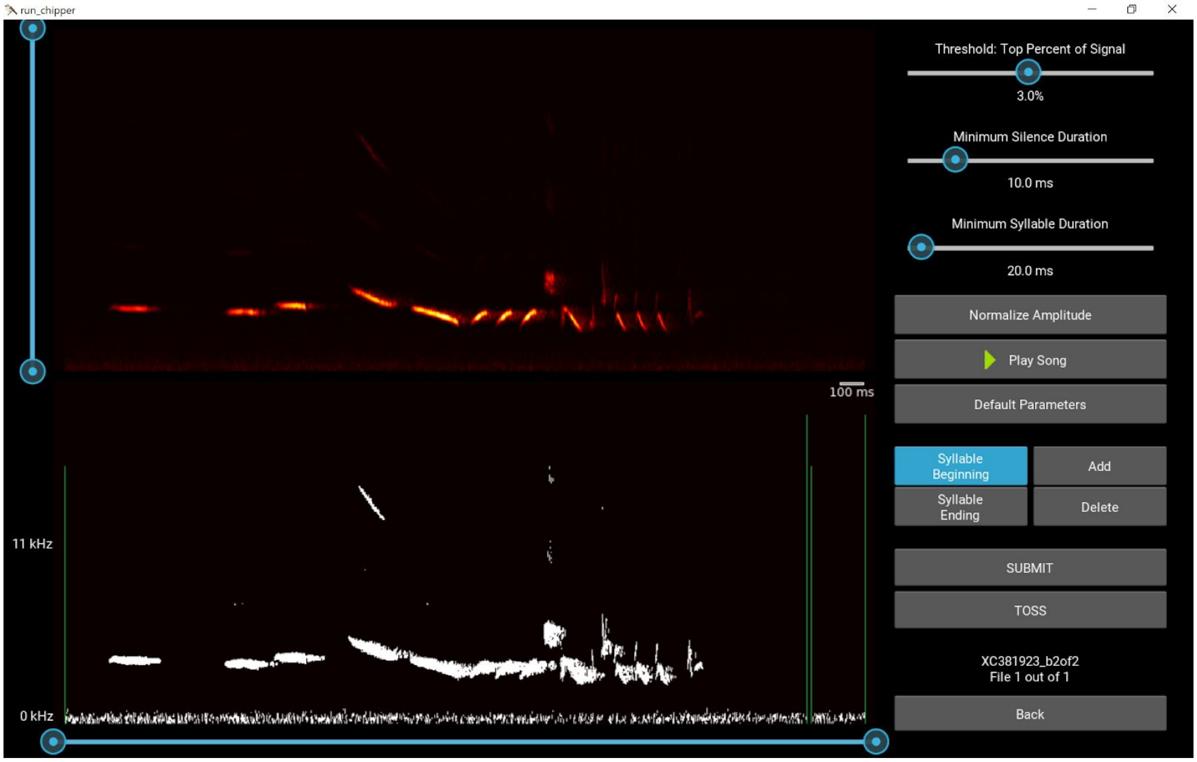

5. Adjust the low-pass and high-pass filters.

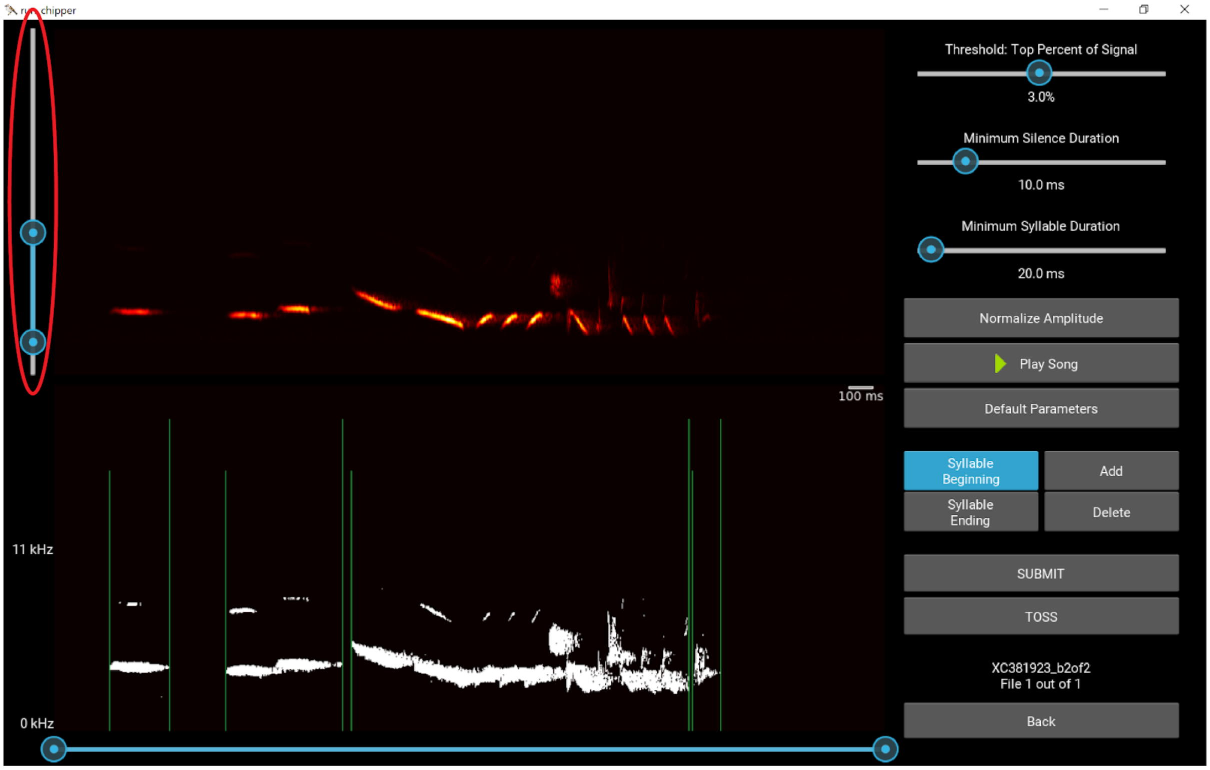


6. Adjust the signal threshold to reduce noise. (You may come back and tweak this again.)

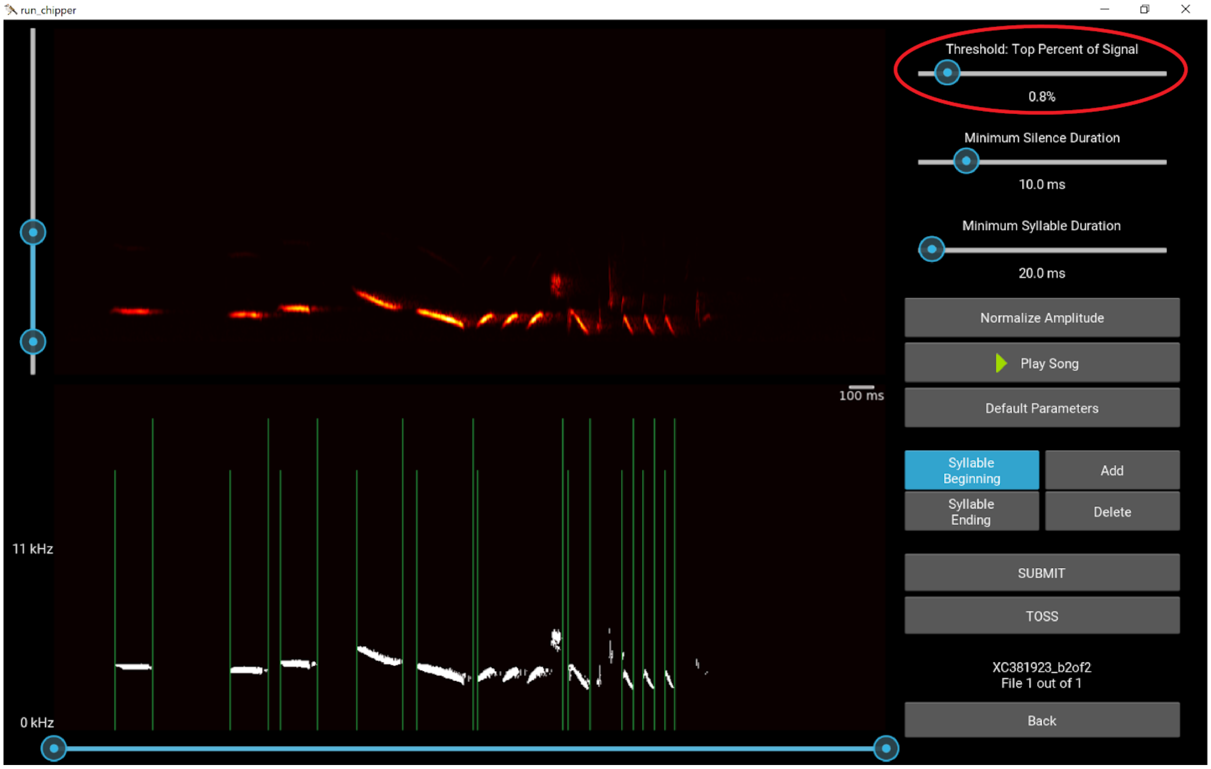

7. Select Normalize Amplitude as some of the small syllables (especially the one on the right end) are fading away with the reduction in signal threshold.

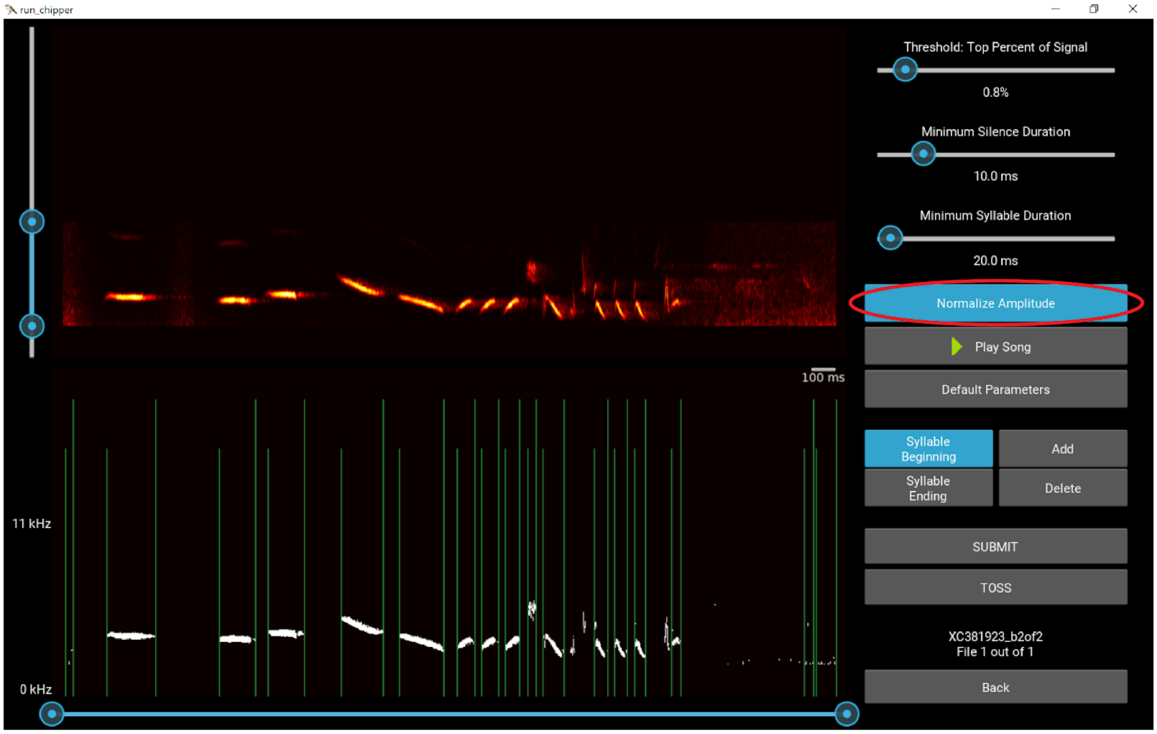


8. Adjust minimum silence duration. This both got rid of some of the onsets and offsets around noise on the ends of the song as well as some of the noise at the end of syllables that appears to be an echo.

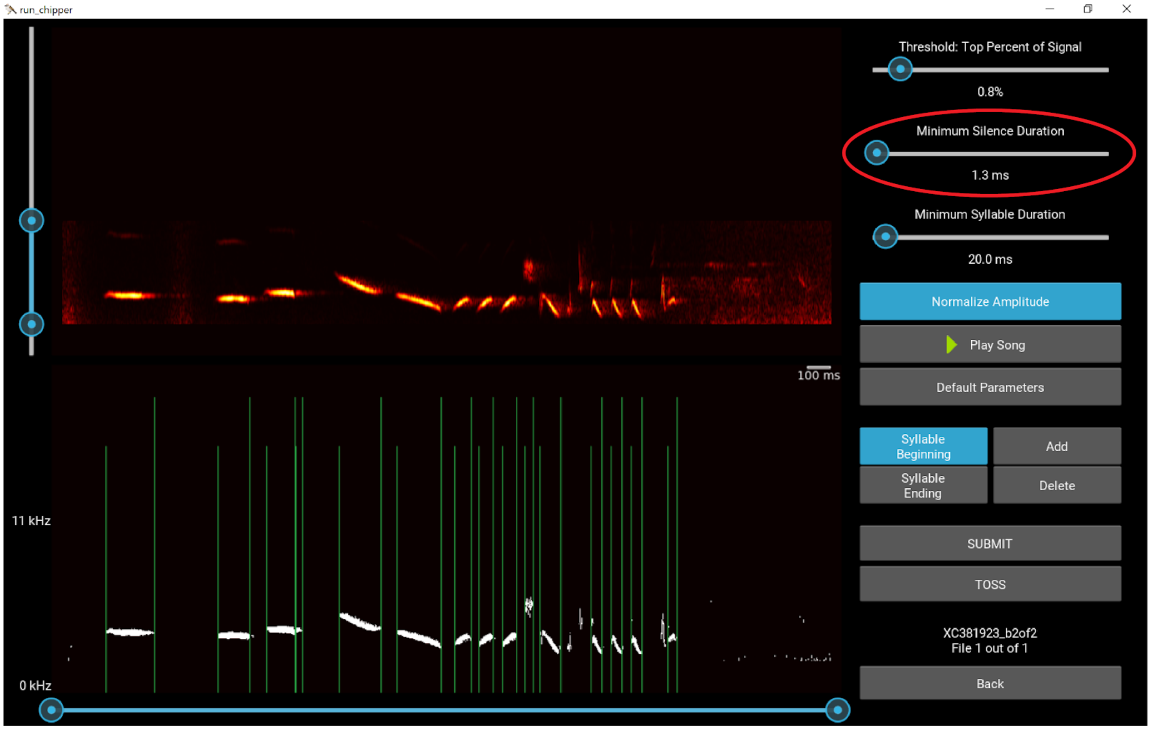

9. Use minimum syllable duration to correctly parse the beginning of the last syllable and the two small syllables in the middle of the song that have no onsets or offsets.

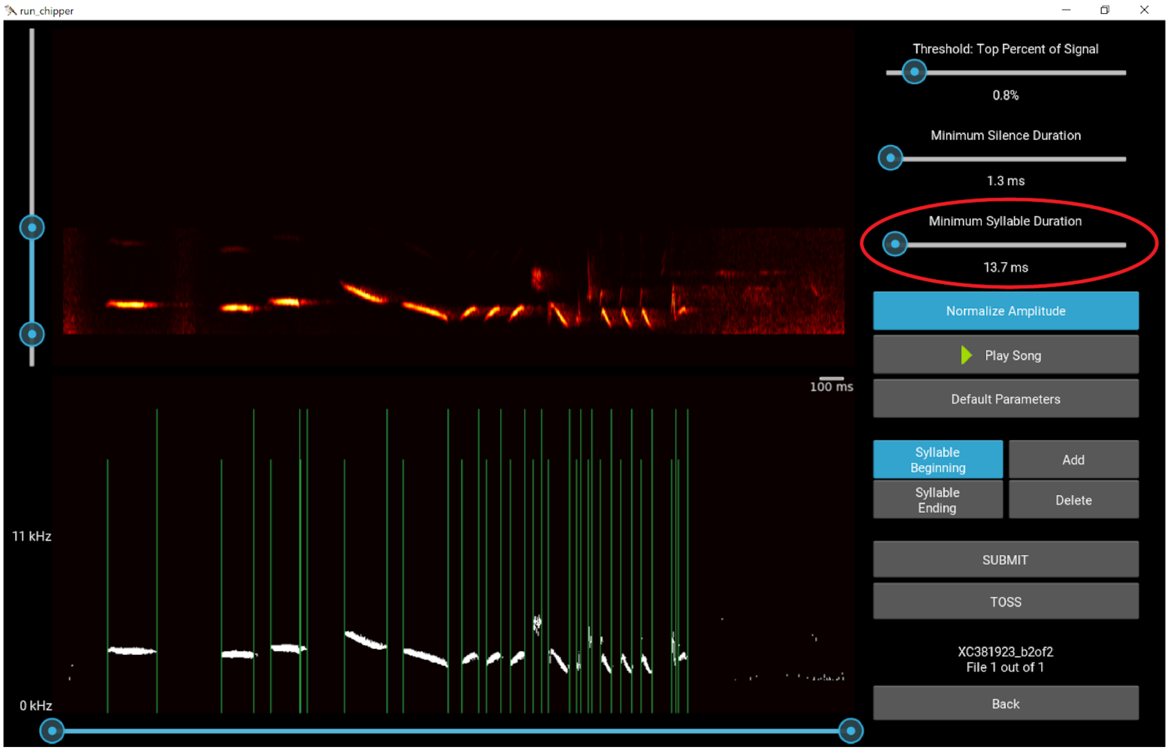


10. Add/Delete any onsets/offsets that are missing or extraneous. Here we delete two syllable beginnings and two syllable endings.

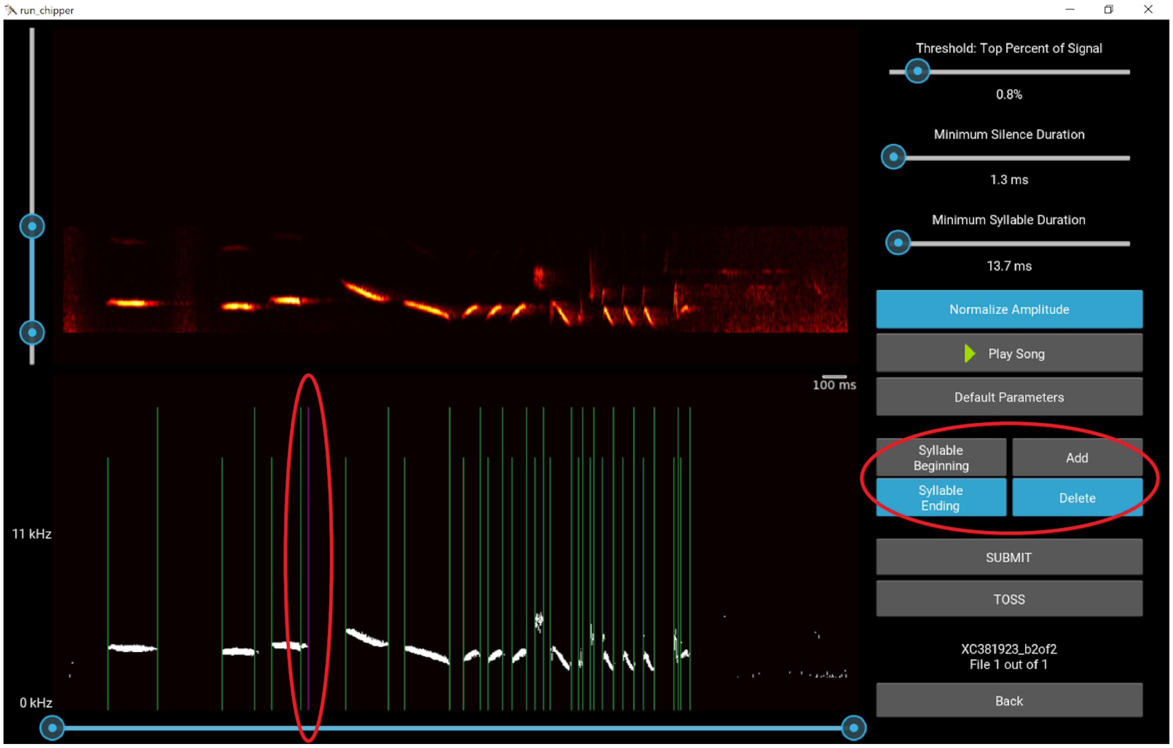

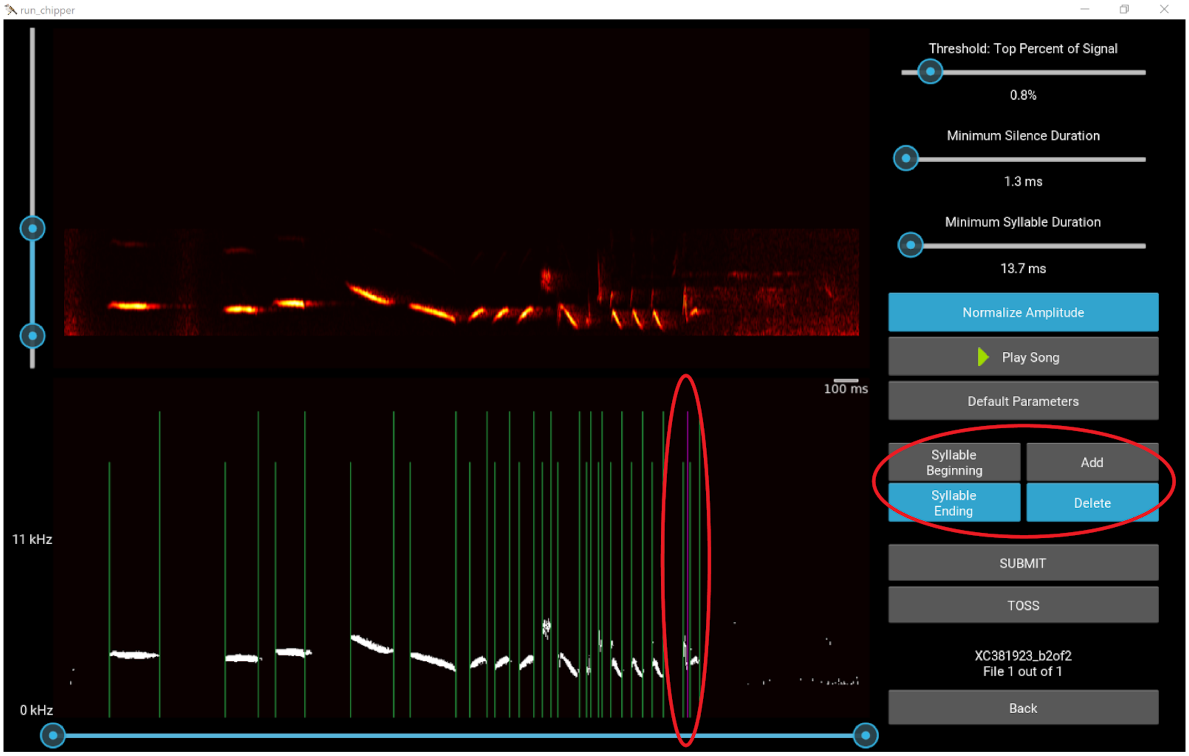
run_chipper
Threshold: Top Percent of Signal
0.8%
Minimum Silence Duration
1.3 ms
Minimum Syllable Duration
13.7 ms
Normalize Amplitude
Play Song
100 ms
Default Parameters
Syllable Beginning
Add
Syllable Ending
Delete
11 kHz
SUBMIT
TOSS
XC381923_b2of2
File 1 out of 1
0 kHz
Back

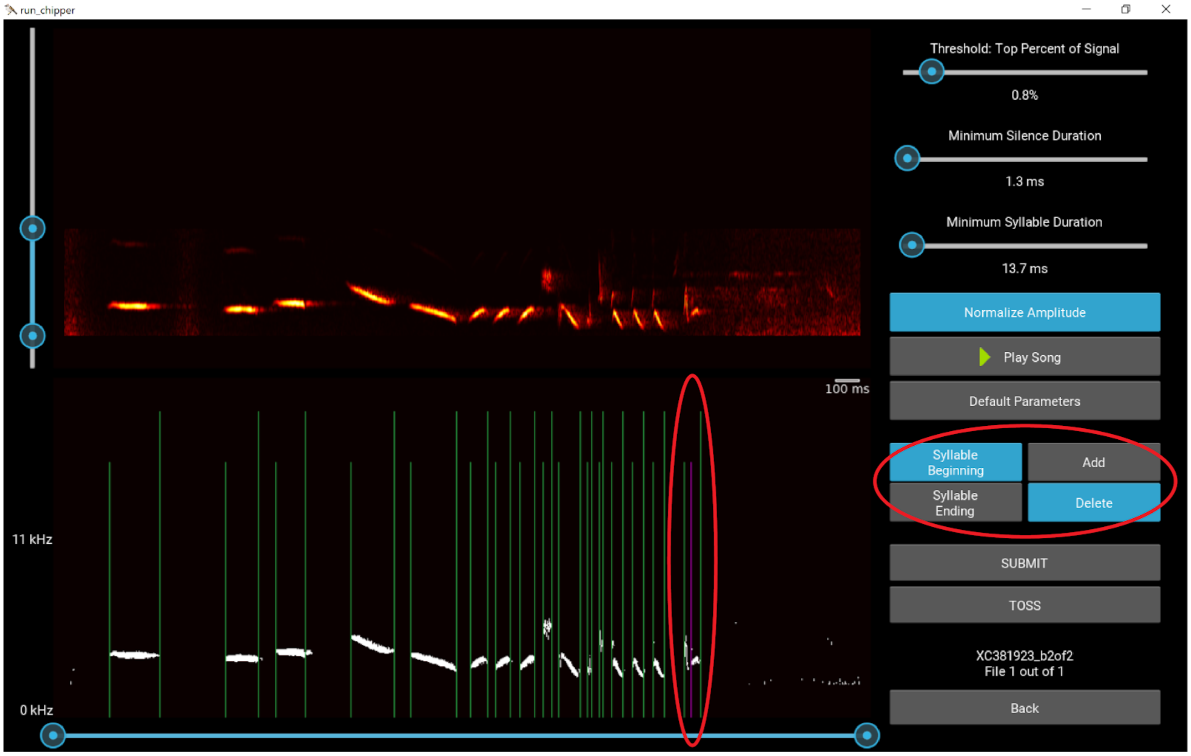
run_chipper
Threshold: Top Percent of Signal
0.8%
Minimum Silence Duration
1.3 ms
Minimum Syllable Duration
13.7 ms
Normalize Amplitude
Play Song
100 ms
Default Parameters
Syllable Beginning
Add
Syllable Ending
Delete
11 kHz
SUBMIT
TOSS
XC381923_b2of2
File 1 out of 1
0 kHz
Back

11. If the noise at the beginning and end of the song were still being parsed as syllables (which is not the case here), you could adjust the bottom slider to crop from the left and right.

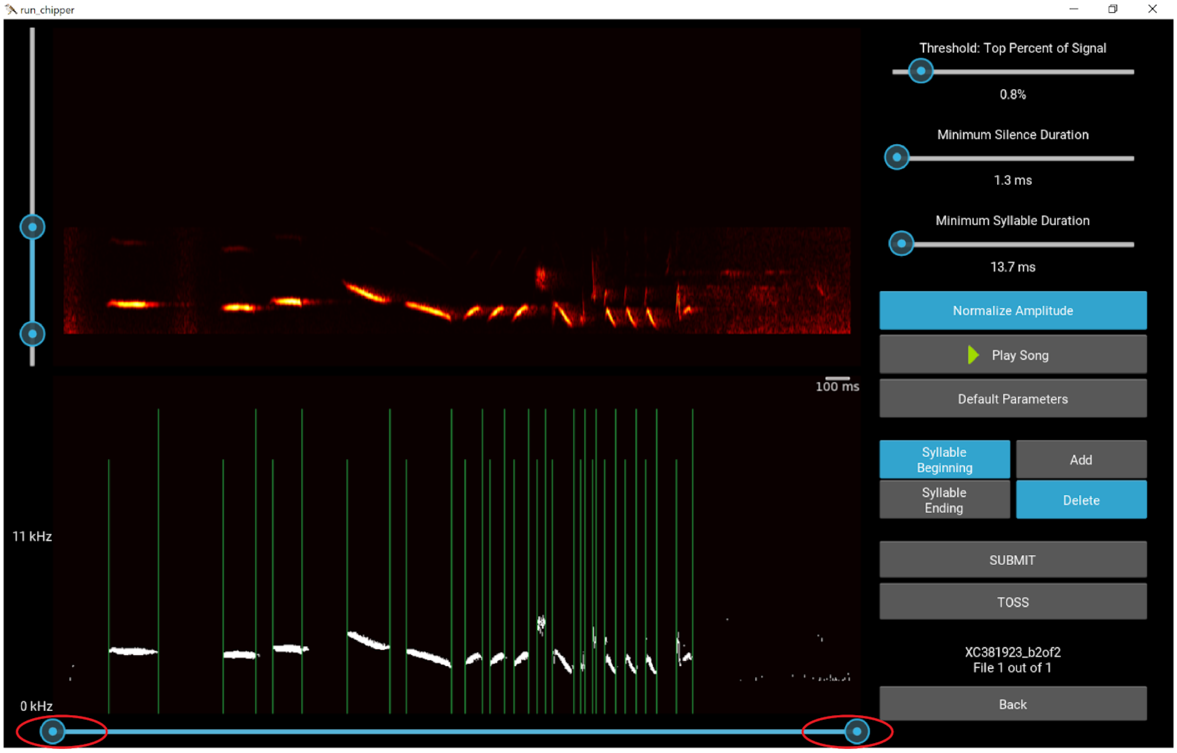


12. Segmentation looks good, select Submit!

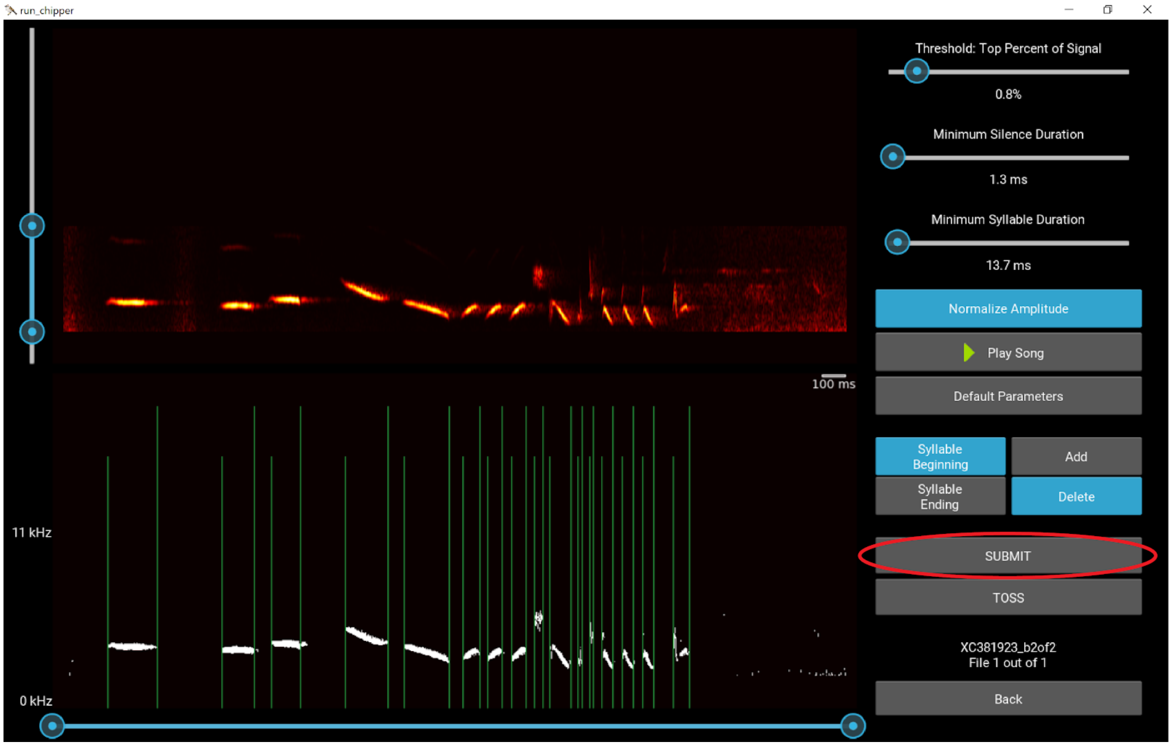

13. Exit Syllable Segmentation and return to the landing page where you can continue to parse another folder of WAV files or can continue with this set in the threshold widgets and analysis.

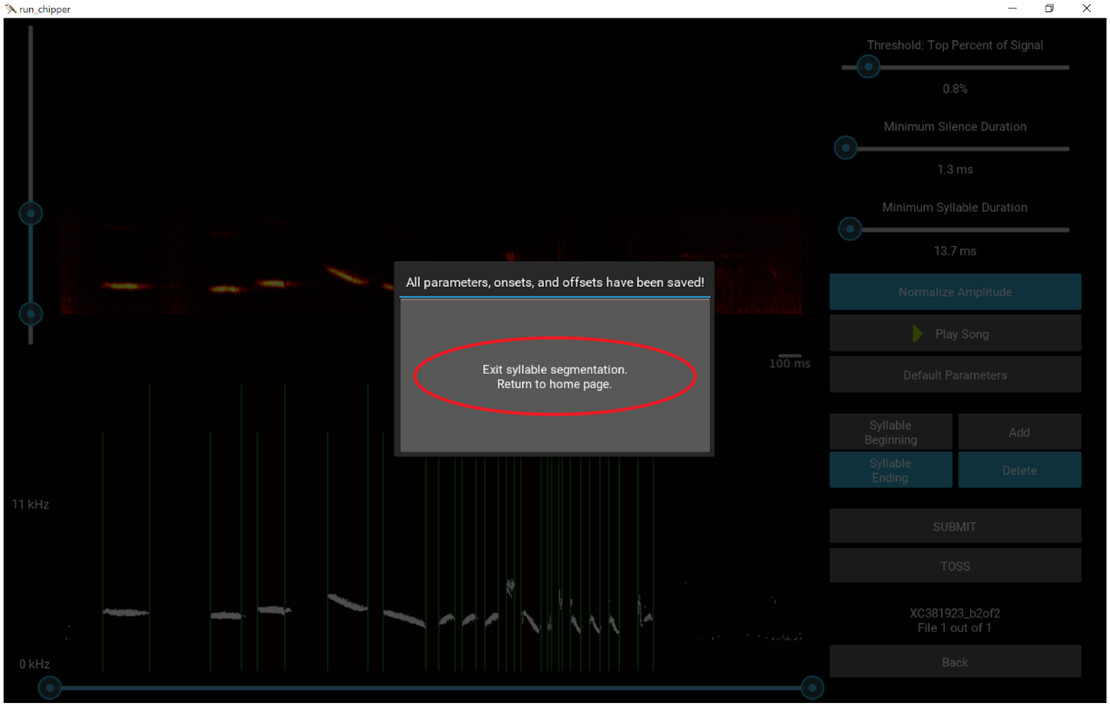


14. The output folder, *SegSyllsOutput_YYYYMMDD_THHMMSS* can now be found in the same folder as the WAV files you ran. This folder contains the four human-readable text file outputs as well as the gzips for each submitted WAV file.

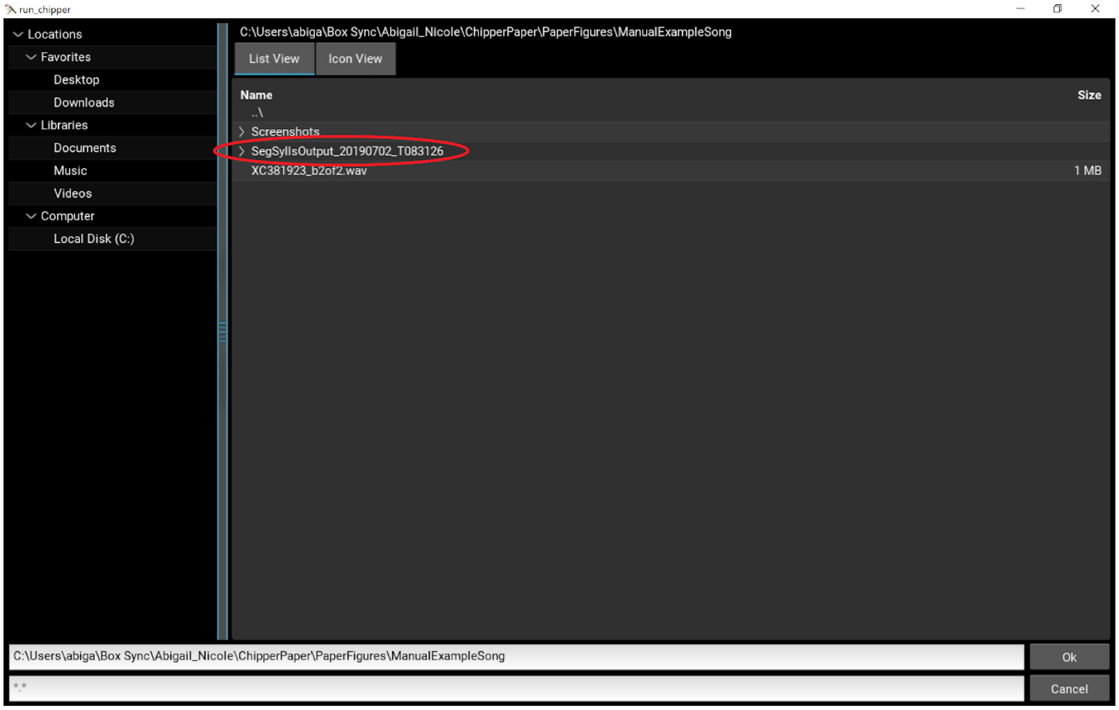

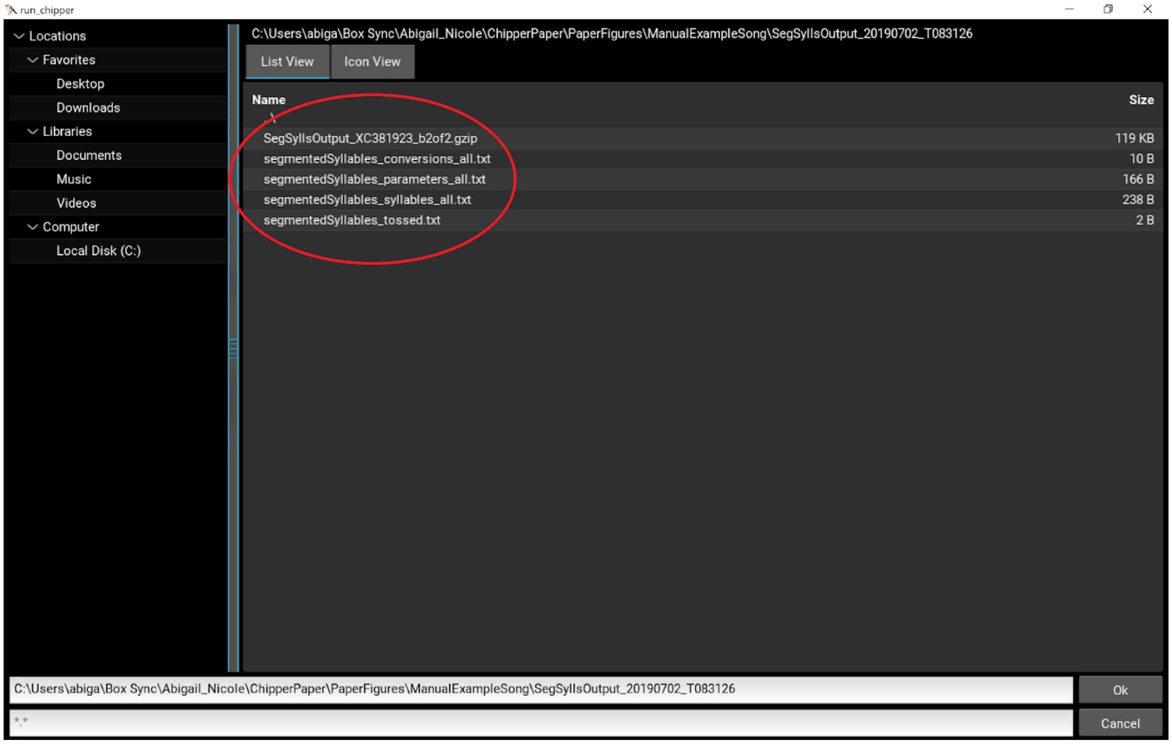

run_chipper
Locations
Favorites
Desktop
Downloads
Libraries
Documents
Music
Videos
Computer
Local Disk (C:)
C:\Users\abiga\Box Sync\Abigail_Nicole\ChipperPaper\PaperFigures\ManualExampleSong\SegSyllsOutput_20190702_T083126
List View
Icon View
Name
Size
SegSyllsOutput_XC381923_b2of2.gzip
119 KB
segmentedSyllables_conversions_all.txt
10 B
segmentedSyllables_parameters_all.txt
166 B
segmentedSyllables_syllables_all.txt
238 B
segmentedSyllables_tossed.txt
2 B
C:\Users\abiga\Box Sync\Abigail_Nicole\ChipperPaper\PaperFigures\ManualExampleSong\SegSyllsOutput_20190702_T083126
Ok
Cancel

**Noise Threshold Widget**

The purpose of the widget is to help you determine a common size threshold for noise for all of your data. Since audio noise often appears on a spectrogram as small pieces of signal, we enable the user to set a size threshold such that sets of connected signal below (or equal to) a certain size are considered noise and not meaningful signal in your recording. Ideally, you have been able to remove all or most noise in the Syllable Segmentation process, such that this widget is primarily functioning to determine separate notes. However, removing small pieces of noise can also be accomplished by setting this threshold.

We recommend you perform this step for a set of songs from the same species. Specifically, you can use a subset of your data (~20 songs) to determine the threshold. You will adjust the threshold for each song until satisfied with the results. A summary of the thresholds used for the sample songs will be given at the end. Then, you will be given the chance to adjust the final threshold to be used in song analysis. You can return to this widget as many times as you wish to visualize the chosen threshold for any songs of interest.

The colors are to help you distinguish separate notes. A note is considered to be a set of connected elements (by edges not corners, e.g. 4-connected) in the binary spectrogram having an area greater than the noise threshold. So, if two notes very close to one another appear separate and are the same color, they are most likely one note. This may be due to the limits of screen resolution. If the area of a note is less than or equal to the noise threshold, it will be considered noise, appearing white in the spectrogram. Noise will not be considered in the analysis calculations.

Below is the example song with the *default* noise threshold. In this case, there only looks to be one syllable with signal that is incorrectly labeled as noise. This signal (circled below) is shown in white, indicating it has few enough spectrogram matrix elements to be considered noise. If we reduce the threshold, smaller notes will be considered signal instead of noise.

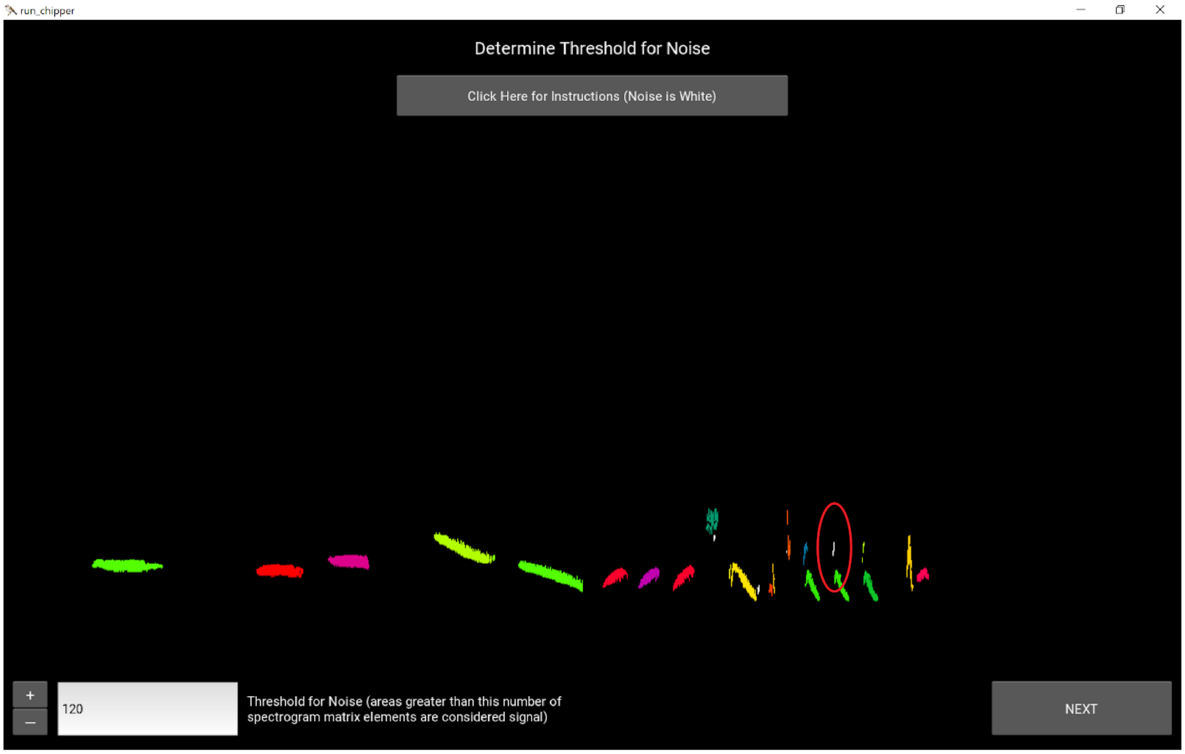


Reducing the noise threshold, the second note of the syllable is now correctly labeled. Here is the example song with the *adjusted* noise threshold.

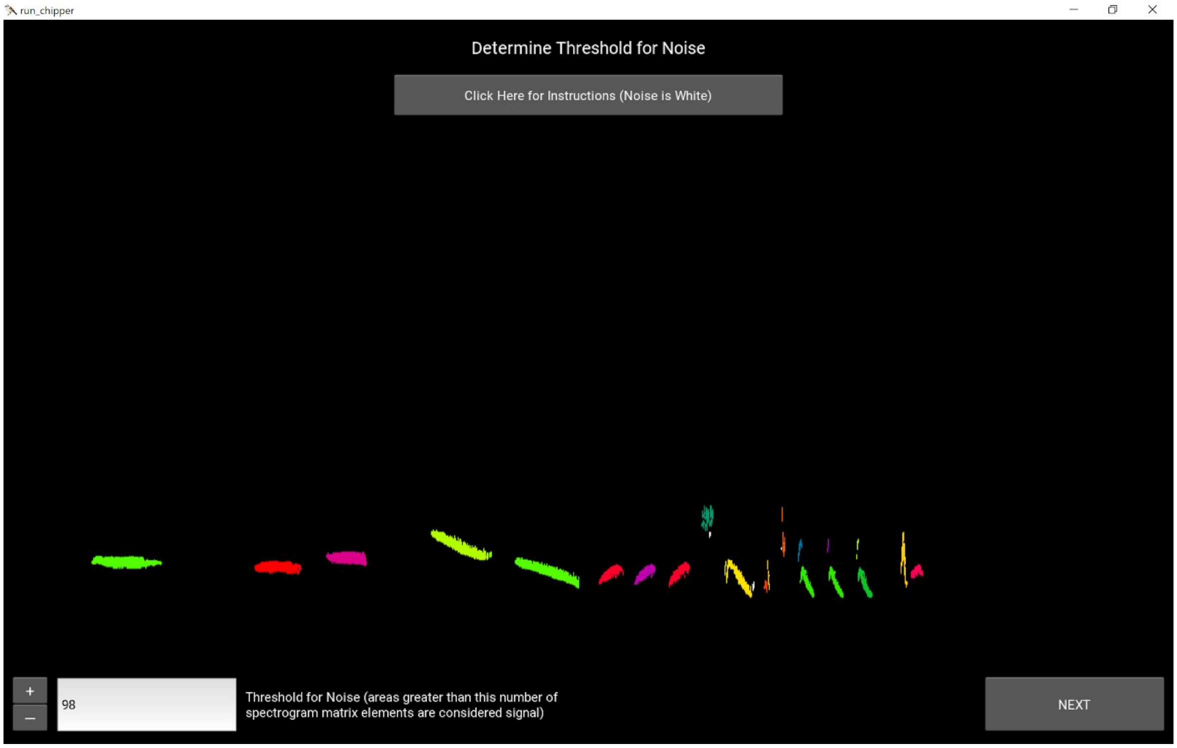

**Syllable Similarity Threshold Widget**

The purpose of the widget is to help you determine a common threshold for syllable similarity for all of your data. When syllables are compared with one another in Chipper, syllables that are more similar to each other than this threshold value (or equal to) will be considered repetitions of the same syllable.

We recommend you perform this step for a set of songs from the same species. Specifically, you can use a subset of your data (~20 songs) to determine the threshold. You will adjust the threshold for each song until satisfied with the results. A summary of the thresholds used for the sample songs will be given at the end. Then, you will be given the chance to adjust the final threshold to be used in song analysis. You can return to this widget as many times as you wish to visualize the segmentation and chosen threshold for any songs of interest.

The colors are to help you distinguish the syntax of the song, which is also written numerically above the spectrogram. Two syllables are considered to be identical if they overlap with an accuracy greater than or equal to the syllable similarity threshold. The syntax is found sequentially, meaning if the second syllable is found to be the same as the first, and the third syllable is found to be the same as the second but not the first, the third will still be the same as both the first and second syllables. To help, the average, minimum, and maximum percent similarity between like syllables is also provided. Note: the minimum can potentially be less than the threshold because syntax is found sequentially.

In the spectrograms shown in this widget, any signal between syllables will be grey and will not be considered in the analysis. Similarly, any noise (determined using the Noise Threshold from the previous step) will be white and will not be considered in the analysis.

Below is the example song with the *default* syllable similarity threshold. In this case, the syntax is close to what we would consider correct for this bird; however, there are a set of three similar syllables at the end that are labeled “12, 13, 12”.

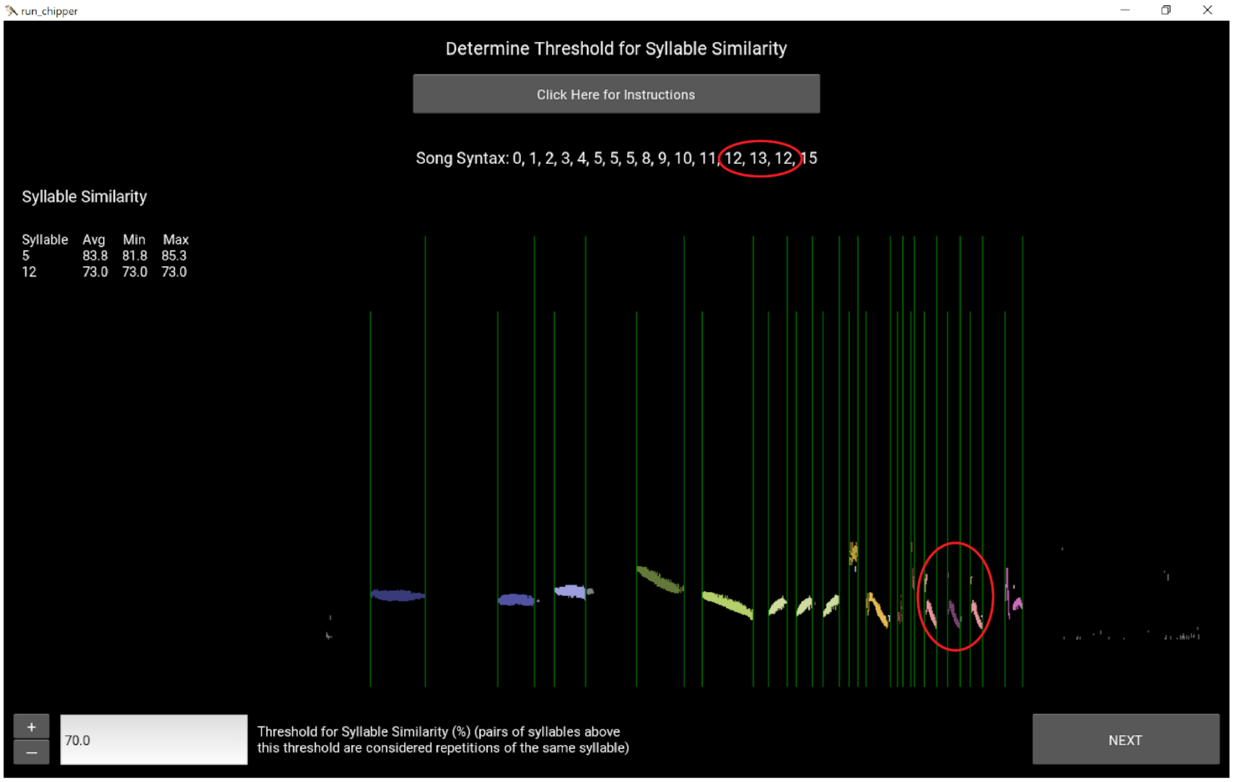


Adjusting the syllable similarity threshold allows us to reach the correct syntax. Here is the example song with the *adjusted* syllable similarity threshold.

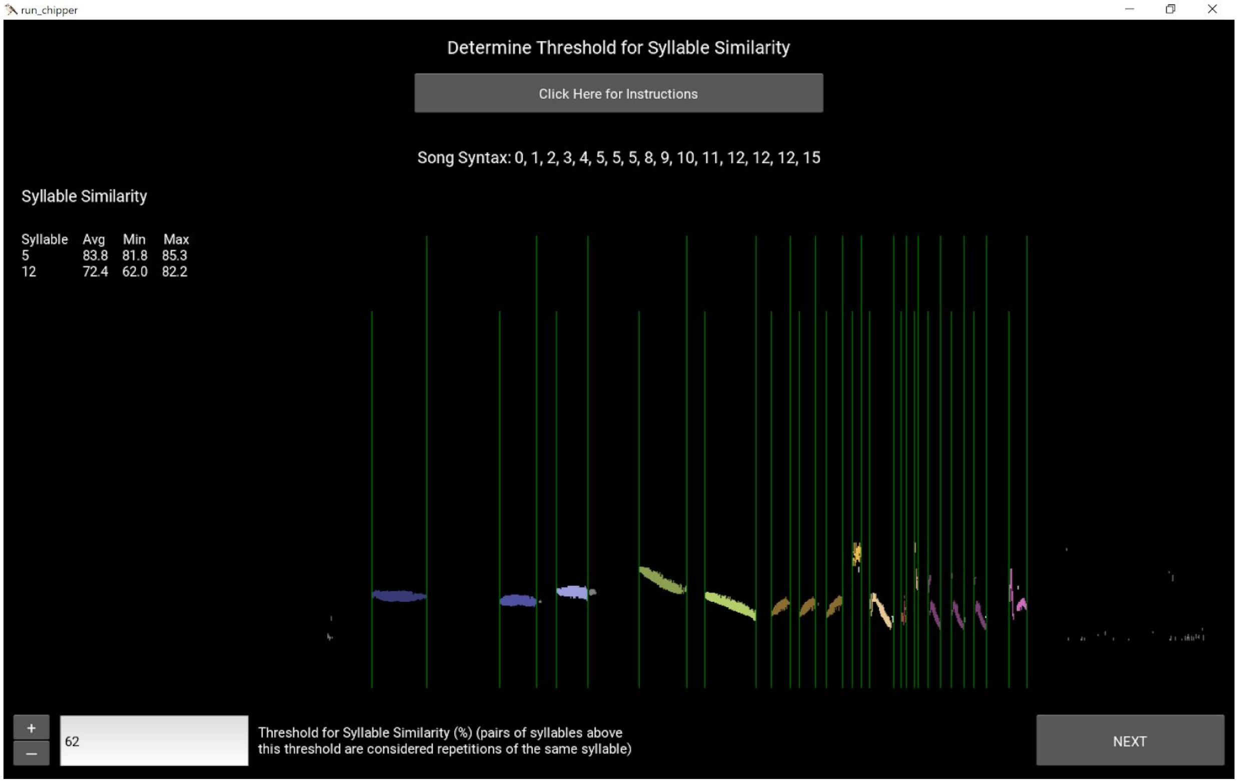

**Running Analysis**

The Noise Threshold and Syllable Similarity Threshold specified on the landing page will be used in Song Analysis. If you have not used the two widgets to determine appropriate thresholds, we recommend doing so before running Song Analysis. Users should be cautious of any note and syntax measurements made in analysis without first finding appropriate thresholds for their data. Similarly, if the user is not concerned with note and syntax information and they are confident they have removed all noise that would artificially raise or lower the frequency measurements, these thresholds can be lowered for analysis.

Starting Song Analysis will first take you to a file explorer to choose a folder of gzip outputs from Syllable Segmentation.

*Note: The gzips do not have to be in their original output folder; the user could have moved them to a new location. If other file types are in the same folder, they will be ignored.*

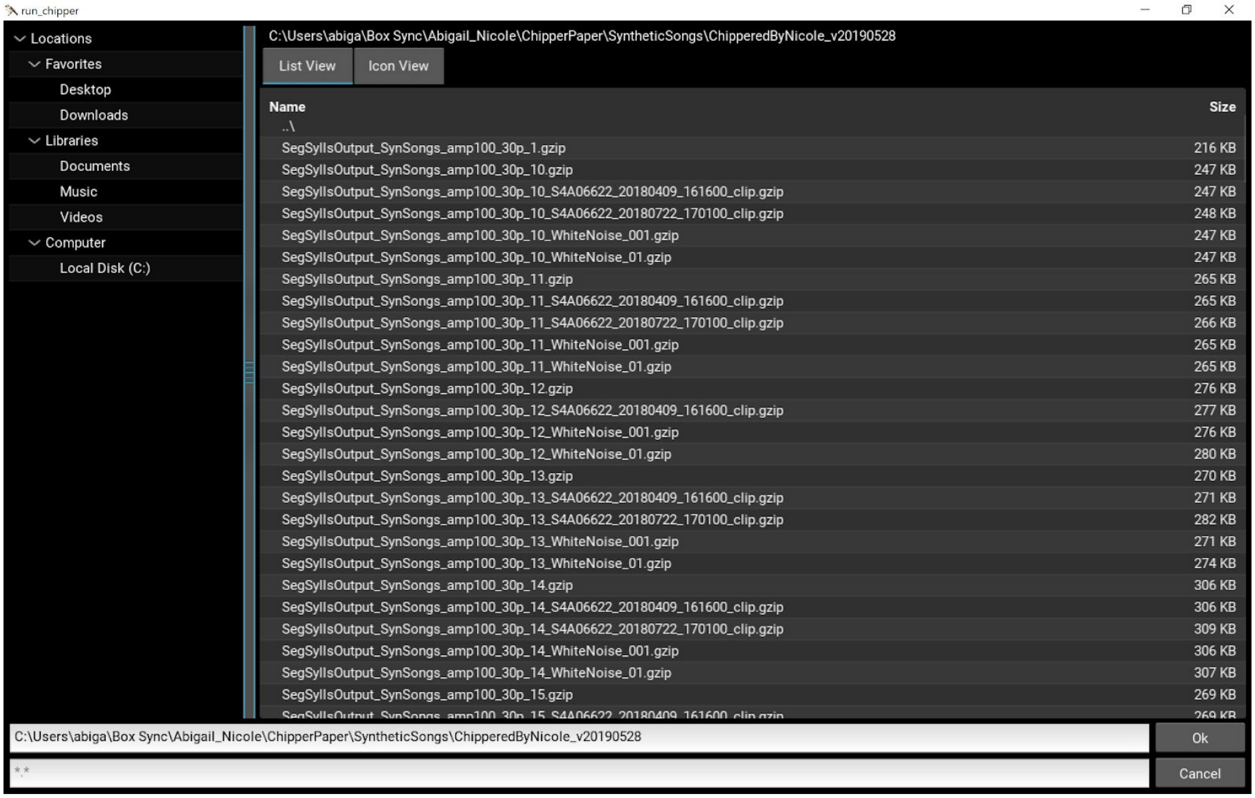


A popup will appear with the number of gzips that will be processed. You can either select "Back" or "Run". When "Run" is selected, a new progress page will appear (see below). The number of files will be incremented as the analysis is completed. Any errors in analysis should throw an exception, printing the error to the analysis page and skipping the current gzip; the error messages will also be logged in a text file named

*AnalysisOutput_YYYYMMDD_THHMMSS_error_log*. While Song Analysis is running, if needed, use the active button to “Cancel Analysis and Exit” Chipper.

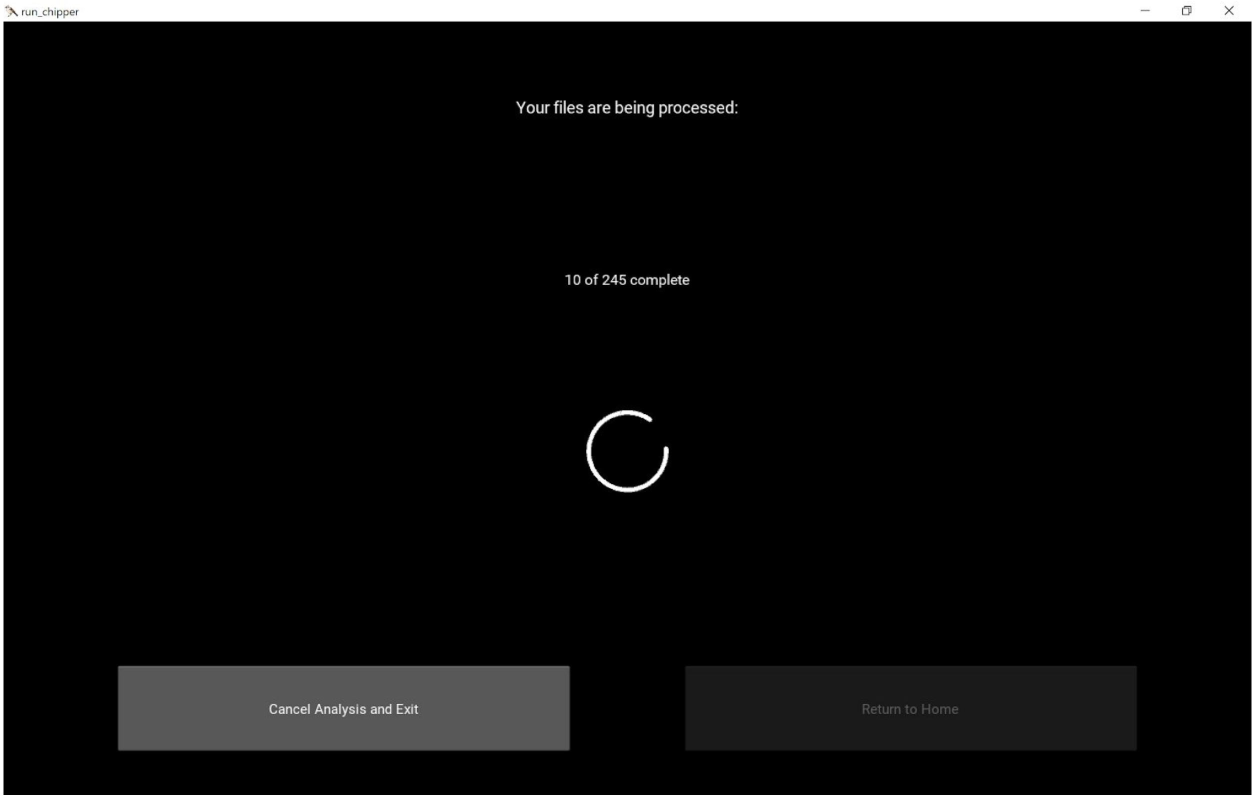


When all gzips are completed, the spinning wheel will no longer be present, and the button “Return to Home” will become active. Two Song Analysis output files can now be found in the folder you analyzed:

1. *AnalysisOutput_YYYYMMDD_THHMMSS_songsylls* with measurements pertaining to the song and syllables.
2. *AnalysisOutput_YYYYMMDD_THHMMSS_notes* with measurements pertaining to the notes. The user should be careful using these measurements, as noisy song files will not have accurate note information due to disconnected signal. In other words, when a syllable is segmented from a noisy file, the signal is likely to be broken up into pieces that are not actually separate notes sung by the bird. Thus, note measurements should be used with caution from field recordings unless the user is confident that syllables are parsed correctly into notes. The user can always visualize the notes using the Noise Threshold widget.

**Analysis Output**

Together the *AnalysisOutput_YYYYMMDD_THHMMSS_songsylls* and *AnalysisOutput_YYYYMMDD_THHMMSS_notes* files include 42 measurements for all gzips run through Song Analysis and the two thresholds submitted by the user for the calculations.

Noise Threshold is used to remove any connected signal with an area less than or equal to the submitted parameter. This is done first, such that all note-related and frequency-related calculations will be affected by this last bit of “cleaning”. Any calculations that exclusively use the onsets and offsets (e.g. syllable duration, silence duration) will not be affected.

Syllable Similarity Threshold is used to determine if two syllables are considered to be repetitions of the same syllable. This affects syllable pattern (syntax) and any measures associated with it—syllable pattern, sequential repetition, syllable stereotypy, and mean and standard deviation of syllable stereotypy.

All Syllable calculations are conducted on the signal between onsets and offsets (i.e. signal that occurs between syllables is not analyzed).

For definitions of each of the measurements see the table below; for more detailed information see [https://github.com/CreanzaLab/chipper/blob/master/chipper/analysis.py](https://github.com/CreanzaLab/chipper/blob/master/chipper/analysis.py).

| Term | Calculation |
|---|---|
| avg_note_duration(ms) | mean(time of note beginning − time of note ending) |
| avg_notes_freq_modulation(Hz) | mean(maximum frequency − minimum frequency for each note) |
| avg_notes_lower_freq(Hz) | mean(minimum frequency of each note) |
| avg_notes_upper_freq(Hz) | mean(maximum frequency of each note) |
| avg_silence_duration(ms) | mean(time of syllable onset − time of previous syllable offset) |
| avg_syllable_duration(ms) | mean(time of syllable offset − time of syllable onset) |
| avg_sylls_freq_modulation(Hz) | mean(maximum frequency − minimum frequency for each syllable) |
| avg_sylls_lower_freq(Hz) | mean(minimum frequency of each syllable) |
| avg_sylls_upper_freq(Hz) | mean(maximum frequency of each syllable) |
| bout_duration(ms) | (time of last syllable offset − time of first syllable onset) |
| largest_note_duration(ms) | max(time of note beginning − time of note ending) |

| | |
|---|---|
| largest_notes_freq_modulation(Hz) | max(maximum frequency − minimum frequency for each note) |
| largest_silence_duration(ms) | max(time of syllable onset − time of previous syllable offset) |
| largest_syllable_duration(ms) | max(time of syllable offset − time of syllable onset) |
| largest_sylls_freq_modulation(Hz) | max(maximum frequency − minimum frequency for each syllable) |
| max_notes_freq(Hz) | max(maximum frequency of each note) |
| max_sylls_freq(Hz) | max(maximum frequency of each syllable) |
| mean_syllable_stereotypy | mean(stereotypy values for each repeated syllable)<br>[see syllable_stereotypy definition below] |
| min_notes_freq(Hz) | min(minimum frequency of each note) |
| min_sylls_freq(Hz) | min(minimum frequency of each syllable) |
| noise_threshold | provided by user (Noise Threshold) [see Noise Threshold Widget]<br><br>Note: Clusters of signal (4-connected elements of the spectrogram) that have an area larger than this threshold are considered notes, and those less than or equal to this threshold are considered noise and removed from analysis. |
| num_notes | number of 4-connected elements of the spectrogram with an area greater than the noise threshold |
| num_notes_per_syll | (total number of notes)/(total number of syllables) |
| num_syllable_per_bout_duration(1/ms) | (number of syllables)/(song duration) |
| num_syllables | number of syllable onsets in a song |
| num_syllables_per_num_unique | (number of syllable onsets in a song)/(unique values in syllable pattern) |
| num_unique_syllables | number of unique values in syllable pattern |
| overall_notes_freq_range(Hz) | max(maximum frequency of each note) − min(minimum frequency of each note) |
| overall_sylls_freq_range(Hz) | max(maximum frequency of each syllable) − min(minimum frequency of each syllable) |
| sequential_repetition | number of syllables that are followed by the same syllable/(number of syllables - 1) |
| smallest_note_duration(ms) | min(time of note beginning − time of note ending) |

| | |
|---|---|
| smallest_notes_freq_modulation(Hz) | min(maximum frequency − minimum frequency for each note) |
| smallest_silence_duration(ms) | min(time of syllable onset − time of previous syllable offset) |
| smallest_syllable_duration(ms) | min(time of syllable offset − time of syllable onset) |
| smallest_sylls_freq_modulation(Hz) | min(maximum frequency − minimum frequency for each syllable) |
| stdev_note_duration(ms) | standard deviation(time of note beginning − time of note ending) |
| stdev_notes_freq_modulation(Hz) | standard deviation(maximum frequency − minimum frequency for each note) |
| stdev_silence_duration(ms) | standard deviation(time of syllable onset − time of previous syllable offset) |
| stdev_syllable_duration(ms) | standard deviation(time of syllable offset − time of syllable onset) |
| stdev_syllable_stereotypy | standard deviation(stereotypy values for each repeated syllable)<br>[see syllable_stereotypy definition below] |
| stdev_sylls_freq_modulation(Hz) | standard deviation(maximum frequency − minimum frequency for each syllable) |
| syll_correlation_threshold | provided by user (Syllable Similarity Threshold) [see Syllable Similarity widget]<br><br>Note: The percent similarity between any pair of syllables is defined as maximum(cross-correlation between each pair of syllables)/maximum(autocorrelation of each of the compared syllables) × 100. If this percent similarity is greater than or equal to the syll_correlation_threshold, the two syllables are considered the same. |
| syllable_pattern | list of the syllables in the order that they are sung, where each unique syllable is assigned a number (i.e. the song syntax) |
| syllable_stereotypy | list of the mean(pairwise percent similarities) for each repeated syllable, where percent similarity is the maximum(cross-correlation between each pair of syllables)/maximum(autocorrelation of each of the compared syllables) × 100 |
